# Accounting for errors in data improves timing in single-cell cancer evolution

**DOI:** 10.1101/2021.03.17.435906

**Authors:** Kylie Chen, Jiří C. Moravec, Alex Gavryushkin, David Welch, Alexei J. Drummond

## Abstract

Single-cell sequencing provides a new way to explore the evolutionary history of cells. Compared to traditional bulk sequencing, where a population of heterogeneous cells is pooled to form a single observation, single-cell sequencing isolates and amplifies genetic material from individual cells, thereby preserving the information about the origin of the sequences. However, single-cell data is more error-prone than bulk sequencing data due to the limited genomic material available per cell. Here, we present error and mutation models for evolutionary inference of single-cell data within a mature and extensible Bayesian framework, BEAST2. Our framework enables integration with biologically informative models such as relaxed molecular clocks and population dynamic models. Our simulations show that modeling errors increase the accuracy of relative divergence times and substitution parameters. We reconstruct the phylogenetic history of a colorectal cancer patient and a healthy patient from single-cell DNA sequencing data. We find that the estimated times of terminal splitting events are shifted forward in time compared to models which ignore errors. We observed that not accounting for errors can overestimate the phylogenetic diversity in single-cell DNA sequencing data. We estimate that 30-50% of the apparent diversity can be attributed to error. Our work enables a full Bayesian approach capable of accounting for errors in the data within the integrative Bayesian software framework BEAST2.

## Introduction

The growth of cancer cells can be viewed as an evolutionary process where mutations accumulate along cell lineages over time. Within each cell, single nucleotide variants (SNVs) act as markers for the evolutionary process. By sampling and sequencing cells, we can reconstruct the possible evolutionary histories of these cell lineages. This can provide insight into the timing of events and modes of evolution.

Currently, there are two main methods for obtaining genomic sequences, bulk sequencing, and single-cell sequencing. Bulk sequencing data is traditionally used in genomic studies. By pooling the genetic material from many cells to form a single observation, greater coverage and thus genetic signal is retained. However, in the context of cancer phylogenetics, the analysis of bulk data poses challenges. Firstly the intermixing of tumor and normal cells affects the genomic signal. Secondly, the pooled sample may be heterogeneous and thus contain a mixture of different genomic variants (Dagogo-Jack and Shaw, 2018; de Bruin et al., 2014; Liu et al., 2018).

In contrast, single-cell sequencing isolates and amplifies the genetic material within a single cell (Kuipers et al., 2017a). The isolation step alleviates the mixture problem. However, errors are more problematic for single-cell sequencing due to insufficient coverage caused by the limited amount of genetic material. The main sources of errors in single-cell sequencing include: cell doublets, where two cells are sequenced as one by mistake; allelic dropout (ADO), where one of the alleles fails to be amplified; and sequencing error, where a base is erroneously read as a different base by the sequencing machine (Kuipers et al., 2017a; Woodworth et al., 2017; Lähnemann et al., 2020). Error models proposed to address these issues include models based on false positives and false negatives (Ross and Markowetz, 2016; Jahn et al., 2016; Zafar et al., 2017, 2019), models of allelic dropout and sequencing errors (Kozlov et al., 2022), and models of read count errors (Satas et al., 2020).

To enable easy integration with molecular clock and phylogeography models that are commonly used in other areas of phylogenetics (Meijer et al., 2012; Malmstrøm et al., 2016; Kearns et al., 2018) we implemented two error models within a mature Bayesian evolutionary framework, BEAST2 (Bouckaert et al., 2019). Our motivation is to enable inference and quantify the uncertainty of both the evolutionary history and model parameters for single-cell phylogenetics. Our paper implements: (i) a model for false positive and false negative errors (Ross and Markowetz, 2016; Jahn et al., 2016; Zafar et al., 2017, 2019) and (ii) a model for ADO and sequencing errors (Kozlov et al., 2022).

We show that our implementation is well-calibrated (Dawid, 1982) and demonstrate these models on real and simulated singlecell DNA data. Our simulation studies show that not accounting for errors leads to inaccurate estimation of timing and substitution parameters when data is error-prone. Our results suggest that using a model that is not error-aware can significantly overestimate the number of substitutions and hence the evolutionary time scale. Analysis of empirical single-cell datasets suggests 30-50% of the phylogenetic diversity can be attributed to errors. Moreover, we show error models are feasible on real datasets, with no additional runtime costs compared to the equivalent non-error version of these models. Finally, we should note that these methods, while developed with cancer analysis in mind, are also applicable to non-cancer single-cell phylogenetics such as somatic cell evolution.

## Related work

For bulk sequencing, there are many tools that estimate the clonal compositions in each bulk sample (Popic et al., 2015; Miura et al., 2018; Jiang et al., 2016) and infer their clonal history (Heide et al., 2019; Cooper et al., 2015; Alves et al., 2019). These clone inference tools are most applicable to bulk sequencing samples that contain a mixture of clones, and phylogenetic reconstruction is performed on the identified clones. However, with the recent availability of single-cell technology, variations between cells can be studied more directly (Schwartz and Schäffer, 2017). This has led to the development of tools for single-cell phylogenetics.

As errors present a key challenge to the analysis of single-cell data, there is a need for models that account for errors introduced during the sequencing process, missing data, coverage discrepancies (Lee et al., 2020), and the ability to quantify uncertainty (Lähnemann et al., 2020).

Early models are based on false positive and false negative errors where the input is a mutation matrix in binary format as in OncoNEM (Ross and Markowetz, 2016), SCITE (Jahn et al., 2016), SiFit (Zafar et al., 2017), or ternary format as in SiFit (Zafar et al., 2017). OncoNEM (Ross and Markowetz, 2016) is a maximum likelihood (ML) method and uses a heuristic search to optimize the likelihood. SCITE (Jahn et al., 2016) is a Bayesian method that uses Markov chain Monte Carlo (MCMC) to sample the posterior but can also be operated in ML mode. Both OncoNEM and SCITE make the infinite sites assumption where a mutation can occur only once at a site. This assumption may be violated on real data, such as by parallel driver mutations (Tarabichi et al., 2021). Besides this, OncoNEM has been shown to be computationally slow, with low phylogenetic accuracy in the presence of ADO (Kozlov et al., 2022). The infSCITE model (Kuipers et al., 2017b) extends SCITE to account for cell doublet errors and test for the infinite sites assumption. SiFit relaxes the infinite sites assumption and additionally accounts for loss of heterozygosity where a single allele is deleted. As deletion events commonly occur across a large region of the chromosome, this could violate the site independence assumption made by the SiFit model.

SCARLET (Satas et al., 2020) implements a read count model which accounts for false positives and false negatives. This is done by correcting read counts at each site using copy-number variation (CNV) output from another software. Empirical studies have shown ADO is the most significant contributor of errors in single-cell DNA sequencing (Wang and Navin, 2015). CellPhy (Kozlov et al., 2022) explicitly models both ADO and sequencing error on diploid genotypes. Unlike models based on false positives and false negatives where different error types are absorbed into the false positive and false negative parameters, CellPhy is a more realistic model of the errors arising from the sequencing process. Furthermore, CellPhy has been shown to produce the most accurate phylogenetic estimates, followed by SiFit and infSCITE on simulated NGS datasets with ADO, amplification, and doublet errors (Kozlov et al., 2022). Both SCARLET and CellPhy make important advances in using data that is closer to the observed sequencing data than previous methods.

Besides CellPhy, which uses the ML phylogenetic framework RAxML (Stamatakis, 2014), other methods are only available as standalone implementations. The advantage of our work is that it enables easy integration with a wide range of population and clock models. In doing so, making these models available for single-cell phylogenetics. This includes relaxed clock models (Drummond et al., 2006), population growth models such as Bayesian skyline plots (Drummond et al., 2005), and phylogeography models such as structured coalescent (Vaughan et al., 2014) and isolation migration models (Nielsen and Wakeley, 2001).

In this paper, we implement error models for binary SNV data and diploid nucleotide SNV data. The binary model accounts for false positive and false negative errors (Ross and Markowetz, 2016; Jahn et al., 2016; Zafar et al., 2017, 2019). The nucleotide model accounts for ADO and sequencing errors (Kozlov et al., 2022). First, we investigate how errors impact the time scale of evolutionary trees inferred from single-cell data. Then, we perform preliminary analyses on real single-cell data to show error models can be used with population growth and molecular clock models. The next section describes the evolutionary models used in this study.

## Materials and Methods

We implement two sets of models: (i) the binary model, which handles mutation presenceabsence data and (ii) the GT16 model, which handles diploid nucleotide genotypes.

The mutation process is modeled as a substitution process evolving along the branches of a tree *τ*, with mutation rates defined by the substitution rate matrix *Q*. Errors are modeled as a noisy process on tip sequences of the tree, where the true genotype is obfuscated according to error probabilities. To perform inference on data, we sample the posterior distribution of trees and the model parameters using Markov chain Monte Carlo (MCMC).

### Software and Input format

Our software is available at www.github.com/bioDS/beast-phylonco. It accepts input files in Nexus, FASTA, or VCF format via a conversion script available at www.github.com/bioDS/vcf2fasta.

### Binary substitution model

The presence or absence of mutation is represented as a binary state Γ = {1, 0}. The rate matrix *Q* has a single parameter, *λ* which is the rate of back-mutation 1 → 0, relative to a mutation rate of 1.

The elements of the rate matrix *Q* are:

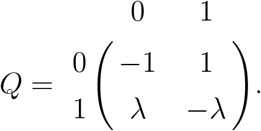

The equilibrium frequencies are:

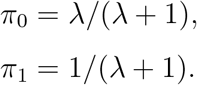

For data sampled at a single time point, the mutation rate is in units of substitutions per site. Data sampled at multiple time points are required to estimate the mutation rate, which typically has units of substitutions per site per year. Alternatively, if we have prior information on the mutation rate, such as from empirical experiments, we can also fix the model’s mutation rate to the empirical value.

### Binary error model

To account for false positive and false negative errors, we implement the binary error model described in (Jahn et al., 2016; Zafar et al., 2017; Ross and Markowetz, 2016). Let *α* be the false positive probability and *β* be the false negative probability. *P* (*x*|*y*) is the conditional error probability of observing noisy data *x*, given that the true state is *y*. For the binary error model, these error probabilities are:

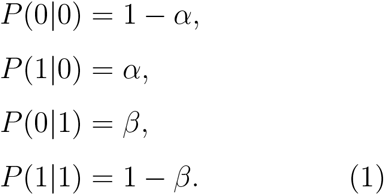

### GT16 substitution model

To model diploid nucleotide sequences, we implement the GT16 substitution and error model described in (Kozlov et al., 2022). The GT16 substitution model is an extension of the four-state general time-reversible nucleotide GTR model (Tavaré et al., 1986) to diploid genotypes: Γ = {*AA, AC, AG, AT, CA, CC, CG, CT, GA, GC, GG, GT, TA, TC, TG, TT*}.

Let *a, b, c, d* be alleles chosen from nucleotides *N* = {*A, C, G, T*} and *r*_*ab*_ be the rate of going from allele *a* to allele *b*.

The elements of the rate matrix *Q* are:

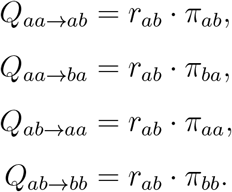

Other non-diagonal entries not listed above have a rate of zero. The diagonals are the sum of the in-going rates:

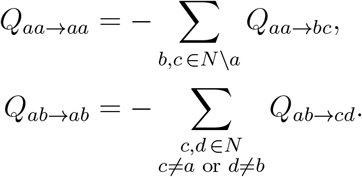

The relative rates of the *Q* matrix are:

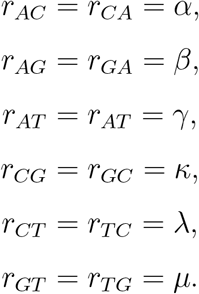

The equilibrium frequencies are: *π* = (*π*_*AA*_, *π*_*AC*_, *π*_*AG*_, *π*_*AT*_, *π*_*CA*_, *π*_*CC*_, *π*_*CG*_, *π*_*CT*_, *π*_*GA*_, *π*_*GC*_, *π*_*GG*_, *π*_*GT*_, *π*_*T A*_, *π*_*T C*_, *π*_*T G*_, *π*_*T T*_).

### >GT16 error model

The GT16 error model for diploid nucleotides described in (Kozlov et al., 2022) accounts for amplification errors and biases in single-cell sequencing. This model has two parameters, the amplification, and sequencing error *ϵ*, and allelic dropout error *δ*. The error probabilities *P* (*x*|*y*) for genotypes with alleles *a, b, c* derived in (Kozlov et al., 2022) are given below:

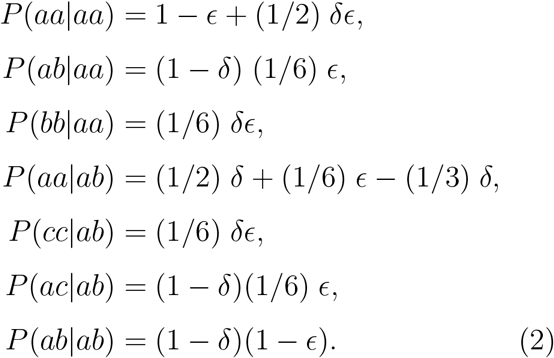

We assume that *P* (*ba*|*aa*) = *P* (*ab*|*aa*) and *P* (*cb*|*ab*) = *P* (*ac*|*ab*). Other combinations not listed above have zero probability. These genotypes can be easily adapted to unphased data by encoding heterozygous states as ambiguities *P* (*ab*^*∗*^) = *P* (*ab*) + *P* (*ba*), where *ab*^*∗*^ represents *ab* without phasing information. Our implementation can handle both phased and unphased data.

### Likelihood calculation

Our input data is a SNV matrix *D* with *n* sites and *m* cells. The cell evolutionary tree *τ* is a rooted binary tree with *m* cells at the leaves, and branch lengths *t*_1_, *t*_2_, … *t*_2(*m*−1)_. These branch lengths are scaled to units of substitutions per site for data sampled at a single time point or years where multiple time points are available. Figure 1 shows an example tree with cells *a, b, c, d* sampled at different time points.

**Figure 1:**
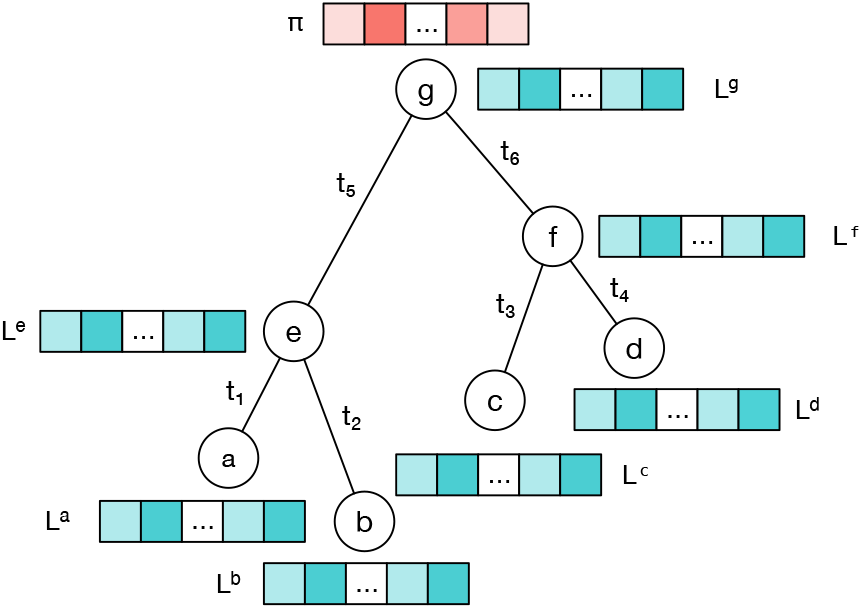
Example of an evolutionary tree with four cells at the leaves *a, b, c, d*.

The likelihood *P* (*D*|*τ, M, θ*) is the conditional probability of observing data *D*, given a tree *τ*, a substitution model *M* with rate matrix *Q* and model parameters *θ*. Assuming each site *i* evolves independently, this likelihood can be written as:

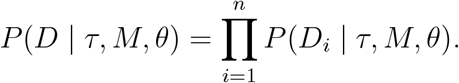

This can be calculated using Felsenstein’s peeling algorithm (Felsenstein, 1981) by recursively traversing the tree. The likelihood at the root node *g, P* (*D*_*i*_|*τ, M, θ*) is calculated by multiplying the equilibrium frequency *π*_*x*_ of genotype *x* at site *i* with its partial likelihood 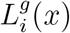 summed over all possible genotypes *x* ∈ Γ:

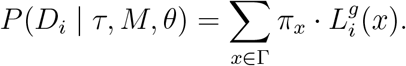

The partial likelihood 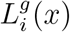 for an internal node *g* at site *i* with child nodes *e* and *f* and corresponding branch lengths *t*_*e*_ and *t*_*f*_ is:

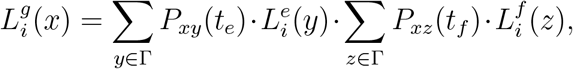

where *P*_*xy*_(*t*) = [*e*^*Qt*^]_*xy*_ is the probabilitiy of going from genotype *x* to genotype *y* after branch length *t* and *Q* is the rate matrix of the substitution model.

Without an error model, the likelihood vector of a leaf node *c* with observed genotype *x* at site *i* is:

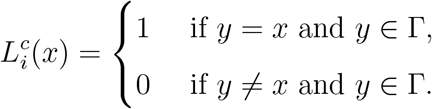

To incorporate errors, we replace the leaf likelihood vectors with the conditional error probabilities *P* (*x*|*y*) in the error model. For example, using an error model the leaf node *c* with observed genotype *x* at site *i* is updated to be:

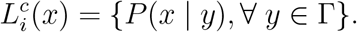

In the binary error model, the leaf likelihood vector 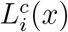 for node *c* is filled using Equation 1 based on its observed genotype *x* at site *i*:

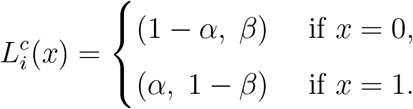

These leaf likelihoods collapse to the non-error version when *α* = 0 and *β* = 0. For brevity, we only fully write out the likelihood of the binary model. The likelihood of leaf nodes for the GT16 error model can be derived similarly using Equation 2 and is available in our implementation.

## Results

### Evaluation on simulated datasets

First, we evaluated our implementations using a well-calibrated study (Dawid, 1982) to test the reliability of the inference when simulating directly from the model. Following the well-calibrated criterion for credible intervals, we expect 95% of the credible interval to cover the true value 95% of the time. Next, we simulated sequences with errors and compared the inference performance with and without modeling the error. Then, we compared the runtime and convergence efficiency of each error model with the baseline non-error substitution model. Lastly, we performed experiments on data simulated with high levels of errors to test the robustness of our methods.

#### Simulation 1: Binary data

We performed a well-calibrated study for the binary model using binary sequences with errors. First, we generated trees using a Yule model, then binary sequences were simulated along the branches of the tree, and errors were applied at the tips. Using the sequence data, we jointly estimate the model parameters, tree topology, and branch times using the binary model.

Simulation parameters: We generated 100 trees with 30 leaves from a Yule model, where each tree has a birthrate drawn from *Normal(µ = 7*.*0, σ = 1*.*0)*. Sequences of length 400 were simulated using the binary model with rate *λ ∼ Lognormal(µ = −1)*, false positive probability *α ∼ Beta(1, 50)* and false negative probability *β ∼ Beta(1, 50)*.

Figure S1 shows the estimates for the model parameters, tree length, and tree height compared to the true simulated values. The estimated 95% highest posterior density (HPD) intervals are shown as bars, where blue indicates the estimate covers the true value, and red indicates otherwise. Our simulations show that the true value of each parameter falls within the estimated 95% HPD interval 91-99% of the time. Figure S2 shows the estimated trees are, on average, 2-5 subtree prune and regraft (SPR) moves away from the true tree.

#### Simulation 2: Binary data error vs. no error

To compare the effects of inference with and without error modeling, we used the data from Simulation 1, then performed inference with and without the binary error model. We compared the coverage for each parameter with and without an error model, i.e., how often the estimated 95% HPD covers the true value. Figure S1 shows the estimated parameters with the binary error model, and Figure S3 shows the estimated parameters without using an error model. Table S1 shows the coverage of each parameter. The coverage of tree length drops from 95% when the error model is used to 39% when no error model is used. Similarly, the coverage of the substitution parameter *λ* drops 91% to 53%. Furthermore, the tree length tends to be overestimated when no error model is used. This suggests that both the tree length and substitution parameters are significantly biased when errors present in the data are not modeled. Figure 2 shows a comparison of the estimated tree length and tree height for these two model configurations. Other parameters such as birthrate and tree height are less biased, with a coverage of 92% and 84% respectively when no error model is used.

**Figure 2:**
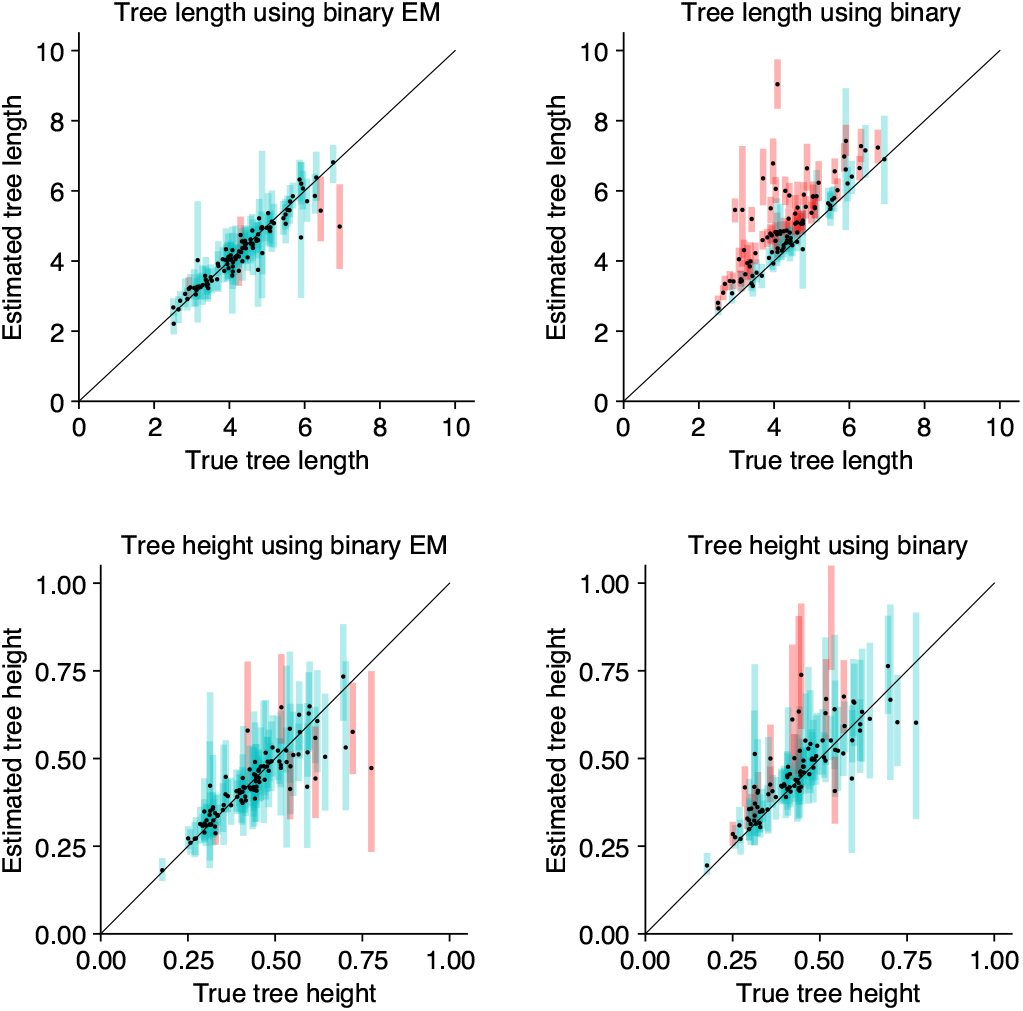
Comparison of branch lengths with and without the binary error model on simulated binary data. Estimated branch lengths using the binary error model (left) and without using the error model (right). Tree length estimates higher than 10, and tree height estimates higher than 1 are truncated on this plot.

#### Simulation 3: Diploid nucleotide phased data

We performed a well-calibrated study for the GT16 model using phased sequence data with errors. First, we simulated trees using a coalescent model. Sequences were simulated down branches of the tree using the GT16 substitution model, and then errors were applied at the tips. Using these sequences as input, we estimated the tree and model parameters using the GT16 model. The priors on the error probabilities were chosen based on experimental studies for allelic dropout (Huang et al., 2015), amplification and sequencing errors (Gawad et al., 2016; Ross et al., 2013).

Simulation parameters: We generated 100 trees with 16 leaves from a coalescent model, where the population size is drawn from *θ ∼ LogNormal(µ = −2*.*0, σ = 1*.*0)*. Sequences of length 200 were simulated using the GT16 model with genotype frequencies *π ∼ Dirichlet(3, 3*, …, *3)*, and relative rates *r ∼ Dirichlet(1, 2, 1, 1, 2, 1)*. Errors were simulated using *E ∼ Beta(α = 2, β = 18)*, and *δ ∼ Beta(α = 1*.*5, β = 4*.*5)*.

Figures S4-S9 show the 95% HPD estimated for each model parameter and tree branch lengths. The true value of each parameter falls within the 95% HPD interval 91-99% of the time. This shows we are able to accurately estimate the substitution, error, population parameters, and branch lengths for phased data. Next, we computed the accuracy of tree topology by comparing the estimated tree with the true tree. Figure S10 shows the average distance from the estimated trees to the true tree. On average, estimated trees are 2-6 SPR moves away from the true tree.

#### Simulation 4: Diploid nucleotide unphased data

We performed a well-calibrated study for the GT16 model using unphased sequencing data. For unphased data, we used the data generated from Simulation 3, with phasing information removed from the sequences. Phasing information was removed by mapping a heterozygous *ab* to both states *ab* and *ba*.

Figures S11-S15 show the estimated model parameters compared to the true simulated values. For each parameter, the estimated 95% HPD interval covers the true value 94-99% of the time, which confirms our implementation is well-calibrated. On average, the estimated trees are 2-6 SPR moves away from the true tree, Figure S16. We note that the sum of the paired heterozygous frequencies (*π*_*ab*_ + *π*_*ba*_) are identifiable, but the individual frequencies (*π*_*ab*_, *π*_*ba*_) are non-identifiable as the data is unphased.

#### Simulation 5: Diploid nucleotide data error vs. no error

To compare the effects of inference with and without error modeling for diploid nucleotide data, we used the data from Simulation 3, then performed inference with and without the GT16 error model.

Table S2 shows the coverage of each parameter with and without an error model. Figures S17-S21 show the estimated model parameters when an error model is not used. We observe a similar trend to Simulation 2, where the tree length and substitution parameters are significantly biased without an error model. Although the tree height estimated without an error model are less biased, the tree lengths are overestimated. These differences in the tree heights and tree lengths are highlighted in Figure 3.

**Figure 3:**
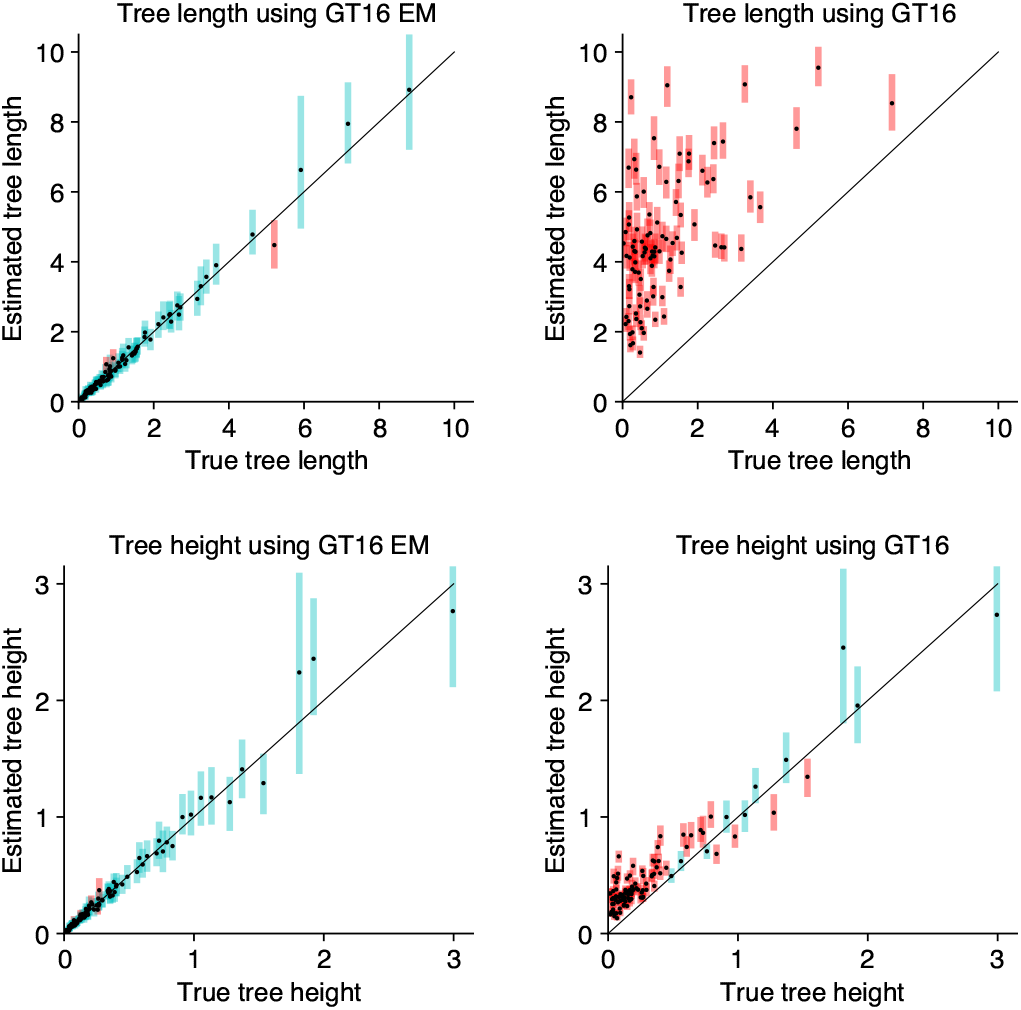
Comparison of branch lengths with and without the GT16 error model on simulated phased nucleotide data. Estimated branch lengths using the GT16 error model (left) and without using the error model (right). Tree length estimates higher than 10, and tree height estimates higher than 3 are truncated on this plot.

#### Simulation 6: Timing experiments

We measured the runtime and convergence of the error model compared with the baseline non-error implementation in our framework. Both error models are comparable in computationally runtime efficiency with their baseline non-error substitution models. Runtime comparisons are shown in supplementary figures S24 - S25. The GT16 model takes approximately an hour to reach convergence on simulated datasets with 20 taxa and 500 sites (convergence is measured as the time till the minimum effective sample size is greater than 200). On a similar-sized dataset, the binary model takes less than five minutes to converge. Timing experiments were done on an Intel Xeon E3-12xx v2 virtual machine with 16 processors at 2.7MHz and 32GB RAM hosted by Nectar Research Cloud.

#### Simulation 7: Performance on data with high levels of error

Lastly, we performed experiments on extended error ranges based on empirical studies (Huang et al., 2015; Gawad et al., 2016; Ross et al., 2013) to test the robustness of our method. We used the same simulation parameters as simulation 1 and simulation 3, but with varying levels of error chosen from an extended range: *α* ∈ [0.001, 0.1], *β* ∈ [0.1, 0.6] for binary data and *δ* ∈ [0.1, 0.8], *E* ∈ [0.001, 0.1] for diploid genotype data. The priors on the error parameters are *α* ∼ *Beta*(1, 20) and *β* ∼ *Beta*(3, 3) for binary data and *δ* ∼ *Beta*(1.5, 4.5) and *E ∼ Beta*(2, 18) for diploid nucleotide data. Our results in the supplementary materials confirm our methods are robust to high levels of error.

### Evaluation on single-cell datasets

We analyzed two public datasets from previously published studies; L86, a colorectal cancer dataset (Leung et al., 2017), and E15, a healthy neurons dataset (Evrony et al., 2015). out an outgroup constraint. The trees estimated using an error model are more treePreprocessed SNVs from CellPhy (Kozlov et al., like than ones estimated without an error 2022) were used for both L86 and E15.

### Colorectal cancer dataset (L86)

L86 contains 86 cells sequenced from a colorectal cancer patient with metastatic spread. The cells were sampled from the primary tumor (colorectal), the secondary metastatic tumor (liver), and matched normal tissue. We used the GT16 model with a relaxed clock to allow for different molecular clock rates in cancer and non-cancer lineages and a coalescent skyline tree prior, which allows changes in population sizes through time.

Model parameters: We used a GT16 substitution model with priors of frequencies *π ∼Dirichlet(3, 3*, …, *3)*, relative rates *r ∼ Dirichlet(1, 2, 1, 1, 2, 1)*, and GT16 error model with allelic dropout *δ ∼ Beta(1*.*5, 4*.*5)* and sequencing error *E ∼ Beta(2, 18)*. A relaxed clock with a Lognormal prior, and Skyline coalescent tree prior with *θ*_1_ *∼ Lognormal(µ = −2*.*3, σ = 1*.*8)*. We performed two independent repeats of the MCMC chains.

We found that the tree height is similar for both error and non-error models, but the relative ages of terminal branches are shorter for the error model. Figure 4 shows the tree length, treeness (Phillips and Penny, 2003; Lanyon, 1988) and gamma statistics (Pybus and Harvey, 2000) of the tree distributions under different experimental setups: with and without error modeling, and with and without an outgroup constraint. The trees estimated using an error model are more treelike than ones estimated without an error model. For the default setup without an outgroup, the 95% HPD estimate for tree length is (4.79, 5.85) with the error model and (7.00, 8.19) without the error model. The error parameters estimates are *δ* ∼ (0.62, 0.66) and *E* ∼ (7 *·* 10^−6^, 1 *·* 10^−3^). The error estimates are comparable to the estimates reported by CellPhy (Kozlov et al., 2022) which are *δ* ∼ 0.63 and *E* ∼ 0.00.

**Figure 4:**
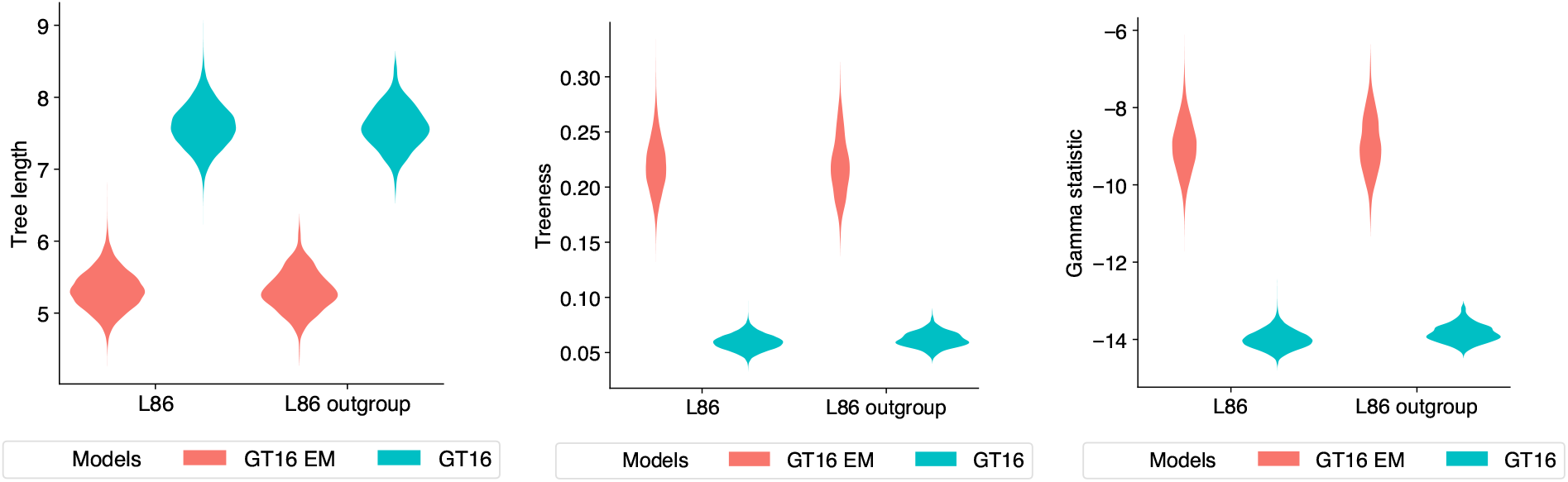
Tree length, treeness and gamma statistics of tree distributions estimated from the L86 dataset. The distributions of each metric is colored by the model used: GT16 error model (red) and GT16 model without error (blue). Two pairs of experiments are shown; L86, which has no tree topology constraints, and L86 outgroup, which has the tree topology constrained to normal cells in the outgroup.

Figure 5 summarizes the estimated tree with the error model (top) compared to without an error model (bottom). The tips of the tree are colored by cell type. The trees show most cells group together by cell type, which suggests there is signal in the data. However, there is some intermixing of metastatic tumor cells inside the primary tumor clade and missorted normal cells as previously identified by (Leung et al., 2017; Kozlov et al., 2022). We also note that the most recent common ancestor (MRCA) of the normal clade is younger than the MRCA of the two tumor clades for both analyses. This is not what we intuitively expected because we believe the normal ancestral cell should be the ancestor of both tumor and normal cells. Although surprising, this observation is in agreement with the trees estimated by ML algorithms in Kozlov et al. (2022). We believe this issue is closely related to the phylogenetic rooting problem for heterogeneous data (Tian and Kubatko, 2017). Methods by Tian and Kubatko (2017), Mai et al. (2017) and Drummond et al. (2006) have provided some partial solutions to the rooting problem, but further research efforts are required to better understand the effects on tree topology.

**Figure 5:**
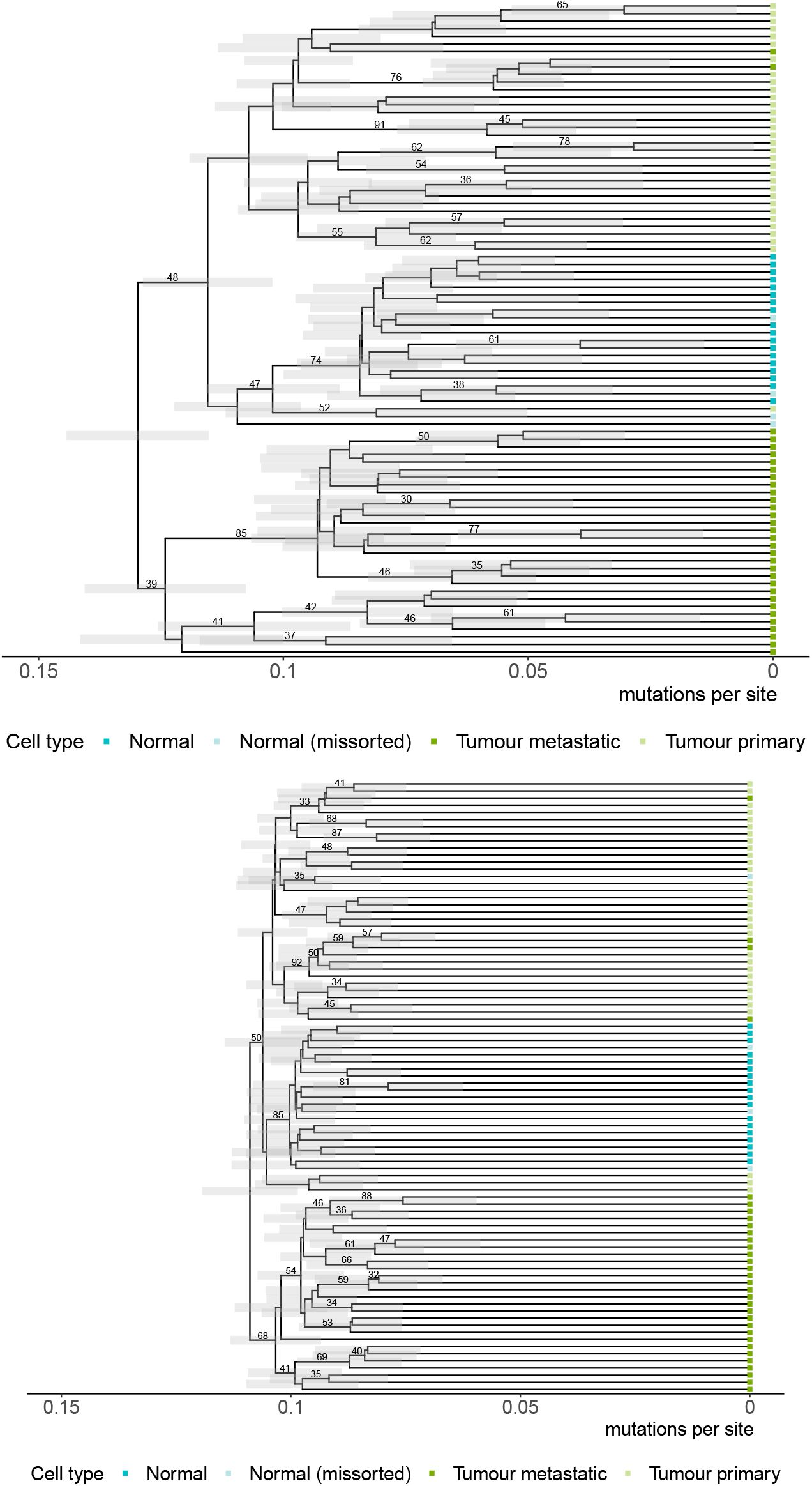
Maximum clade credibility trees for the L86 dataset (colorectal cancer patient) using the GT16 model with an error model (top) and without an error model (bottom). Cells are colored by their cell types: normal cells (dark blue), normal cells missorted during data collection (light blue), the metastatic tumor from the liver (dark green), and primary tumor from the colon (light green). Clades with greater than 30% are labeled.

We also investigated whether constraining the tree topology to have all tumor cells as the ingroup produces different estimates. We repeat the analyses with an outgroup constraint to the tree topology, setting the normal and missorted samples as the outgroup. Figure S22 shows the outgroup constrained summary tree for L86. In this outgroup constrained tree, the age of the MRCA of the normal group is also younger than the MRCA of the primary or metastatic tumors for the error model. The age of the MRCA of the normal group is indistinguishable from the MRCA of the primary or metastatic tumors for the model without error.

### Healthy neurons dataset (E15)

E15 contains 15 neurons and a blood cell taken from the heart region sequenced from a healthy patient.

Model parameters: We used a GT16 substitution with frequencies prior *π ∼ Dirich-let(3, 3*, …, *3)* and relative rates *r ∼ Dirich-let(1, 2, 1, 1, 2, 1)*, GT16 error model with allelic dropout error *δ ∼ Beta(1*.*5, 4*.*5)*, and sequencing error *E ∼ Beta(2, 18)*, with a relaxed clock and Skyline coalescent tree prior with *θ*_1_ *∼ Lognormal(µ = −2*.*3, σ = 1*.*8)*. We performed two independent repeats of the MCMC chains.

We observed that the trees estimated using the error model are more tree-like than ones estimated without an error model, as shown by the tree metrics in figure 6. The 95% HPD estimates of tree length are (1.37, 7.14) with the error model, and (7.60, 13.41) without the error model. We note the tree height for the error model (0.20, 0.80) is lower than that of the non-error model (0.54, 1.01). The estimated interval for the error parameters are *δ* ∼ (0.86, 0.92) and *E* ∼ (0.03, 0.17). To test the sensitivity of the error priors, we reran our experiments with adjusted priors *E, δ ∼ Beta*(1, 10) and *E, δ ∼ Beta*(1, 20). We found the error estimates were similar regardless of these adjustments on the error parameter priors.

**Figure 6:**
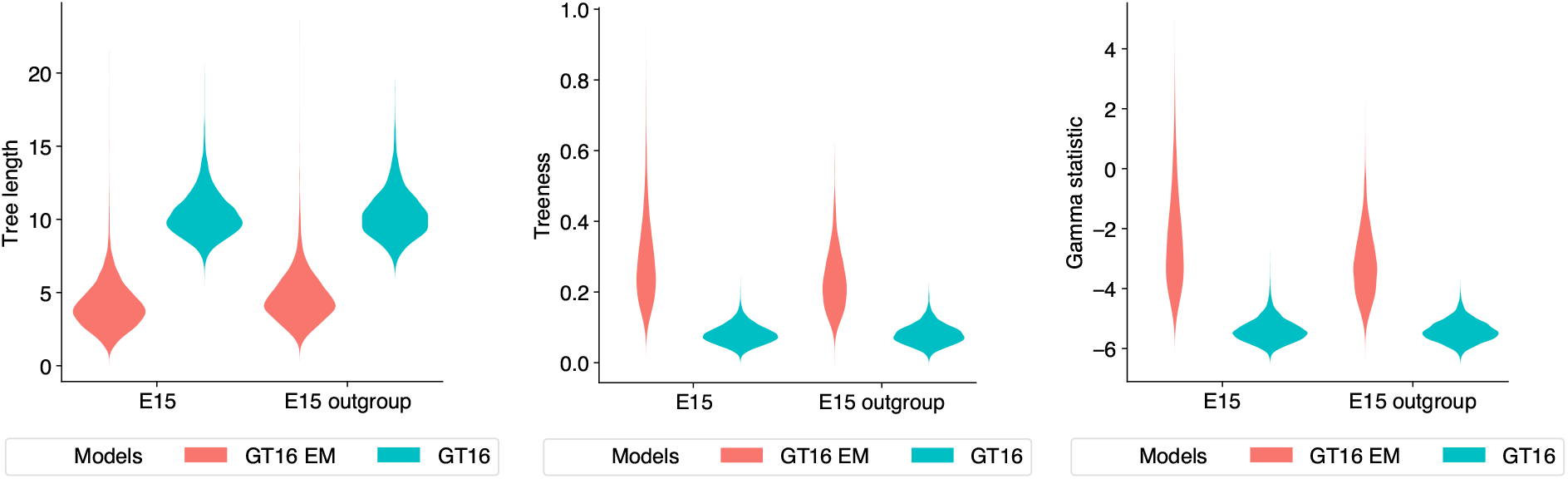
Tree length, treeness and gamma statistics of tree distributions estimated from the E15 dataset. The distributions of each metric is colored by the model used: GT16 error model (red) and GT16 model without error (blue). Two pairs of experiments are shown; E15, which has no tree topology constraints, and E15 outgroup, which has the tree topology constrained with the heart cell as the outgroup.

Figure 7 shows a summary of the estimated trees with the GT16 error model (top) and without an error model (bottom). The tips of the tree are colored by cell types. We expect the blood cell to be placed as an outgroup; however, the estimated trees placed the blood cell inside a clade of neuron cells. To investigate if adding an outgroup constraint to the tree topology can help direct the likelihood in the correct direction, we repeated the analyses with the blood cell as the outgroup. The estimated trees are shown in figure S23. Besides the correct placement of the outgroup enforced by the outgroup constraint, we did not observe any substantial topological discrepancies between the outgroup and non-outgroup analyses.

**Figure 7:**
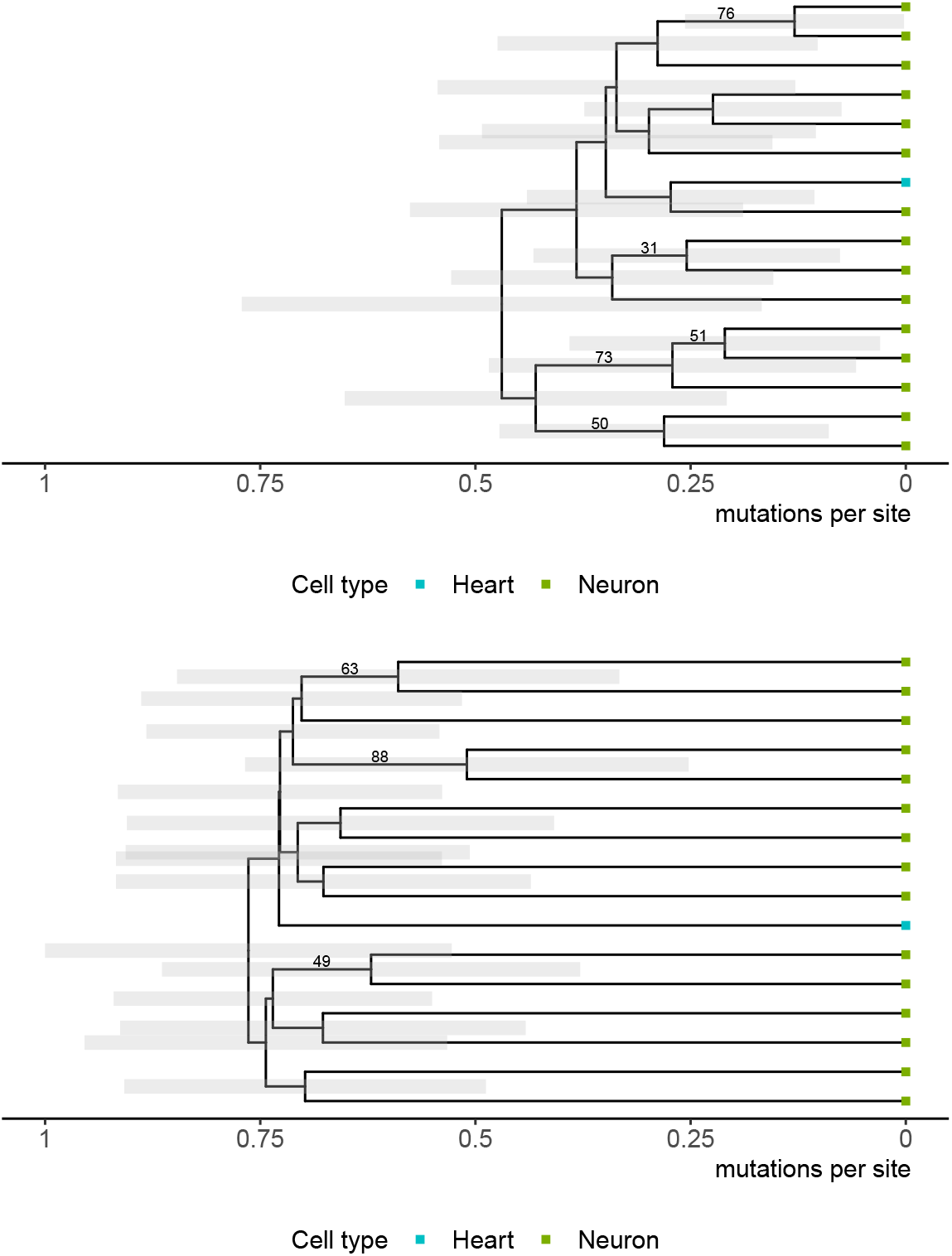
Maximum clade credibility trees for the E15 dataset (healthy patient) using the GT16 model with an error model (top) and without an error model (bottom). Each cell is colored by its cell type: blood cell (blue) and neuron cells (green). The posterior clade support for clades with greater than 30% support are shown on the branches.

## Discussion

We demonstrated that incorporating error parameters can affect the relative ages of singlecell datasets. We showed that models incorporating sequencing error could increase the accuracy of tree branches and model parameters inferred from noisy data. Additionally, we find that using error models is just as fast as the baseline non-error substitution models in our framework. Future work to support multi-threading and add compatibility with the Beagle high-performance library (Ayres et al., 2012) would further increase the computational speed of these models.

From both simulated and real single-cell data, we observed that using an error model tends to shorten the total tree length, as errors explain a portion of the genetic variability within the data. For empirical single-cell data, cells of the same type tend to be placed in the same clade. We believe relaxed clock and local clock models are more suited to heterogeneous data as they allow for changes in mutation rates. The datasets we explored in this paper are sampled at a single time point, so there is no calibration information to allow the mutation rate and time to be disambiguated. Using time sampled data or empirical mutation rate calibrations would improve current analyses and allow node ages to be converted to real time (Drummond et al., 2003, 2002).

Although the effect of filtering strategies in the context of macroevolution shows stringent filtering of sites often leads to worse phylogenetic inference (Tan et al., 2015). The effect of filtering strategies on noisy data such as singlecell phylogenies is yet to be systematically explored. We believe the error parameters in these models can provide increased flexibility, allowing key features of the sequencing and filtering process to be accounted for during evolutionary inference.

Lastly, incorporating cell biology knowledge during method development would improve the biological significance of model assumptions; and improving the interpretability of tree summarization metrics would enable single-cell phylogenies to be examined in more detail.

## Code and data availability

Our software, Phylonco v0.0.6 is available at www.github.com/bioDS/beast-phylonco. Analyses and data are available at www.github. ter Xie for implementation support. We accom/bioDS/beast-phylonco-paper.

This paper uses BEAST v2.6.6 (Bouckaert et al., 2019), BeastLabs v1.9.7, LPhy v1.2.0, and LPhyBeast v0.3.0. The following python packages were used: DendroPy (Sukumaran and Holder, 2010), lxml (Behnel et al., 2005), matplotlib (Hunter, 2007), numpy (Harris et al., 2020), and seaborn (Waskom, 2021). The following R packages were used: ggtree (Yu et al., 2017), ggplot2 (Wickham, 2016), tracerR (Rambaut et al., 2018), treeSimGM (Hagen and Stadler, 2018), treeio (Wang et al., 2020), expm, and ape (Paradis et al., 2019).

## Acknowledgements

We thank Prof. David Posada and Dr. Joao Alves for generously providing us with singlecell datasets and advice. We thank Dr. Remco Bouckaert for helpful discussions and Dr. Walknowledge Nectar Research Cloud and Otago Computing Services for providing the computing resources necessary for this study. AG and AJD were funded by the MBIE Endeavour Fund, ref. U00X1912; AG was supported by Rutherford Discovery Fellowship ref. RDFUOO1702; AJD was supported by the James Cook Research Fellowship from the Royal Society of New Zealand. KC was supported by a University of Auckland Doctoral Scholarship.

