## Supplementary Materials for "Accounting for errors in data improves timing in single-cell cancer evolution"

**This PDF file includes:**

Figures S1 to S31

Tables S1 to S6

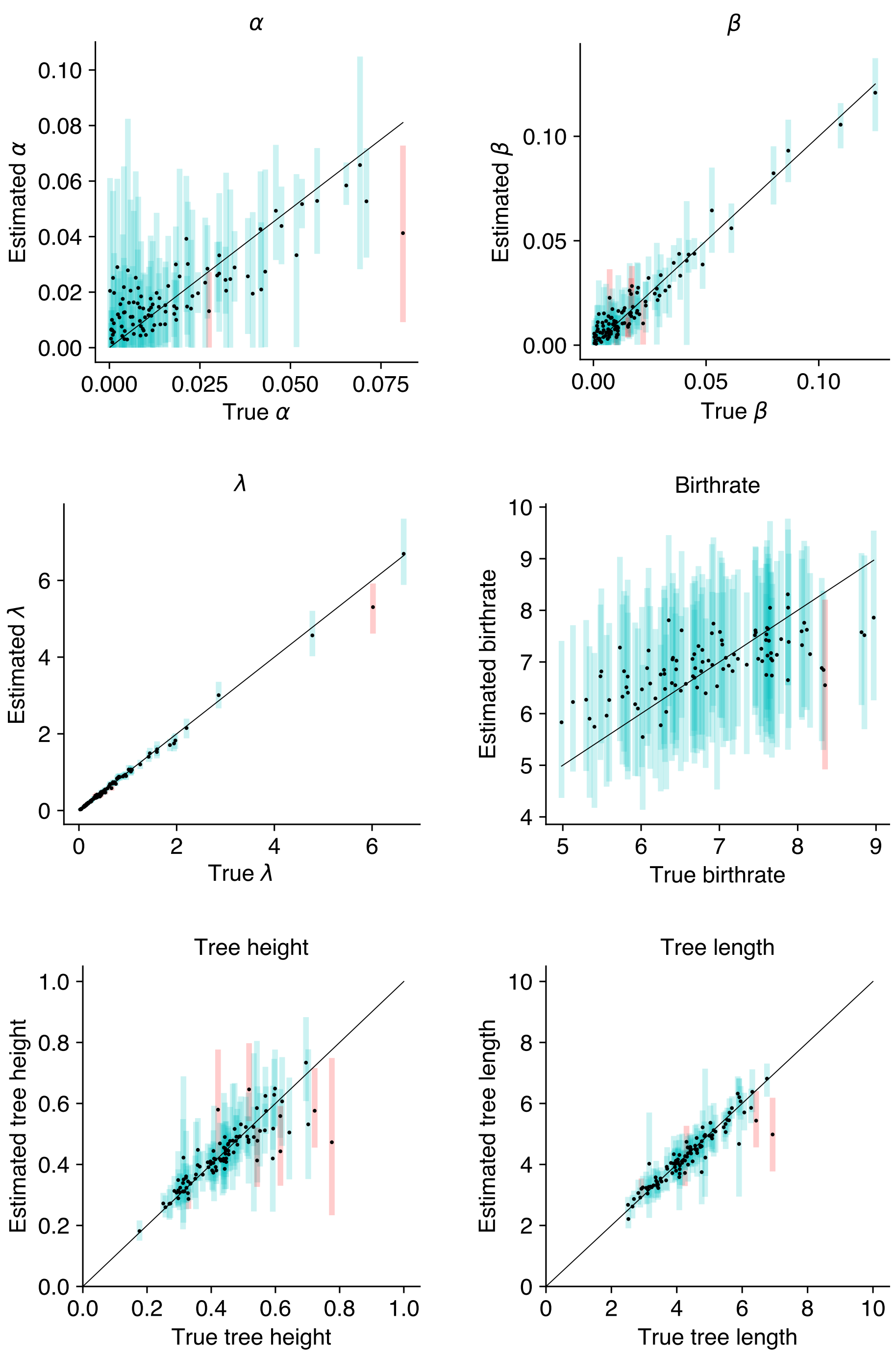

**Figure S1**: Simulation 1: Estimated branch lengths and model parameters on binary data. True vs. estimated parameter values, from left to right, $\lambda$ the relative substitution rate $1 \to 0$ of the binary model, $\alpha$ the false positive rate, $\beta$ the false negative rate, the birthrate of the Yule process, the tree height and the total tree length. Estimated means are shown as points, with true values along the diagonal. The estimated 95% HPD are shown as bars; blue indicates the true value lies within the estimated interval, and red indicates the true value is outside of the estimated interval.

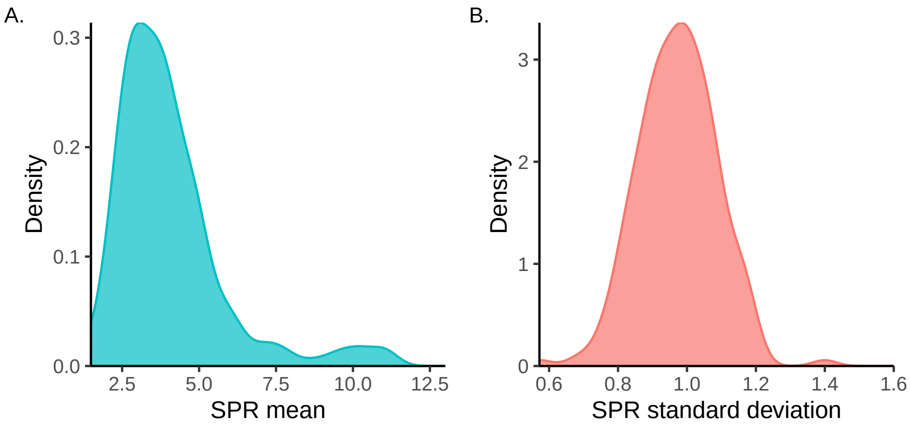

**Figure S2**: Simulation 1: Summary of subtree prune and regraft (SPR) metrics for trees estimated from binary data. From left to right, (A) the mean SPR distance between estimated trees and the true tree, (B) the standard deviation of the SPR distance between estimated trees and the true tree.

**Table S1**: Simulation 2: Coverage comparison for model parameters with and without the binary error model.

| Parameter | 95% HPD coverage with error model | 95% HPD coverage without error model |
|---|---|---|
| $\alpha$ | 98% | - |
| $\beta$ | 93% | - |
| $\lambda$ | 91% | 53% |
| Birthrate | 99% | 92% |
| Tree height | 93% | 84% |
| Tree length | 95% | 39% |

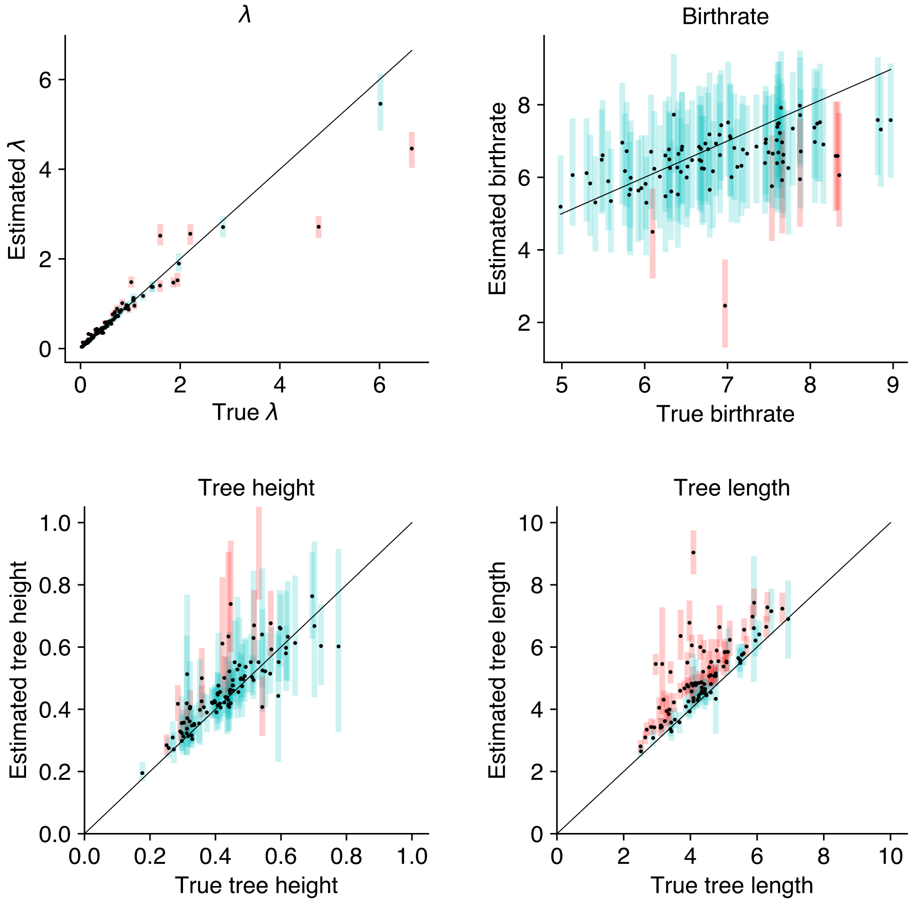

**Figure S3**: Simulation 2: Estimated branch lengths and model parameters on binary data without an error model. True vs. estimated parameter values, from left to right, $\lambda$ the relative substitution rate $1 \to 0$ of the binary model, the birthrate of the Yule process, the tree height and total tree length. Estimated means are shown as points, with true values along the diagonal. The estimated 95% HPD are shown as bars; blue indicates the true value lies within the estimated interval, and red indicates the true value is outside of the estimated interval.

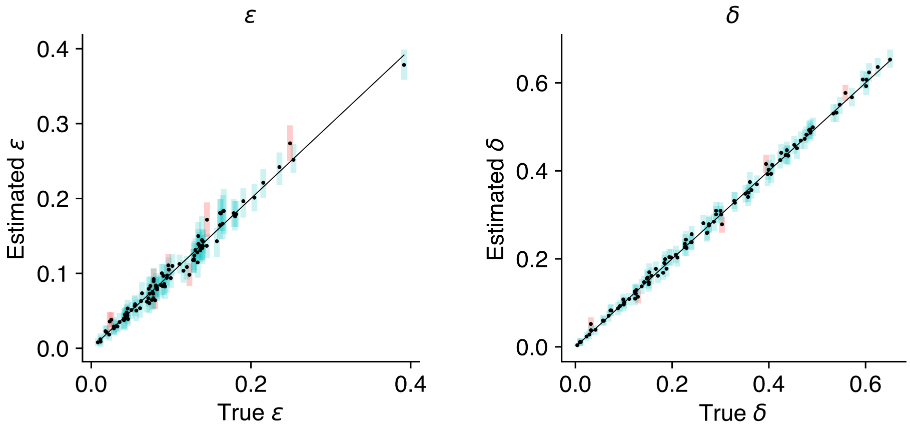

**Figure S4**: Simulation 3: Estimated error parameters for phased diploid nucleotide data. True vs. estimated error probabilities $\epsilon$ and $\delta$ for the GT16 error model using phased data. Estimated means are shown as points, with true values along the diagonal. The estimated 95% HPD are shown as bars; blue indicates the true value falls within the estimated interval, and red indicates the true value is outside of the estimated interval.

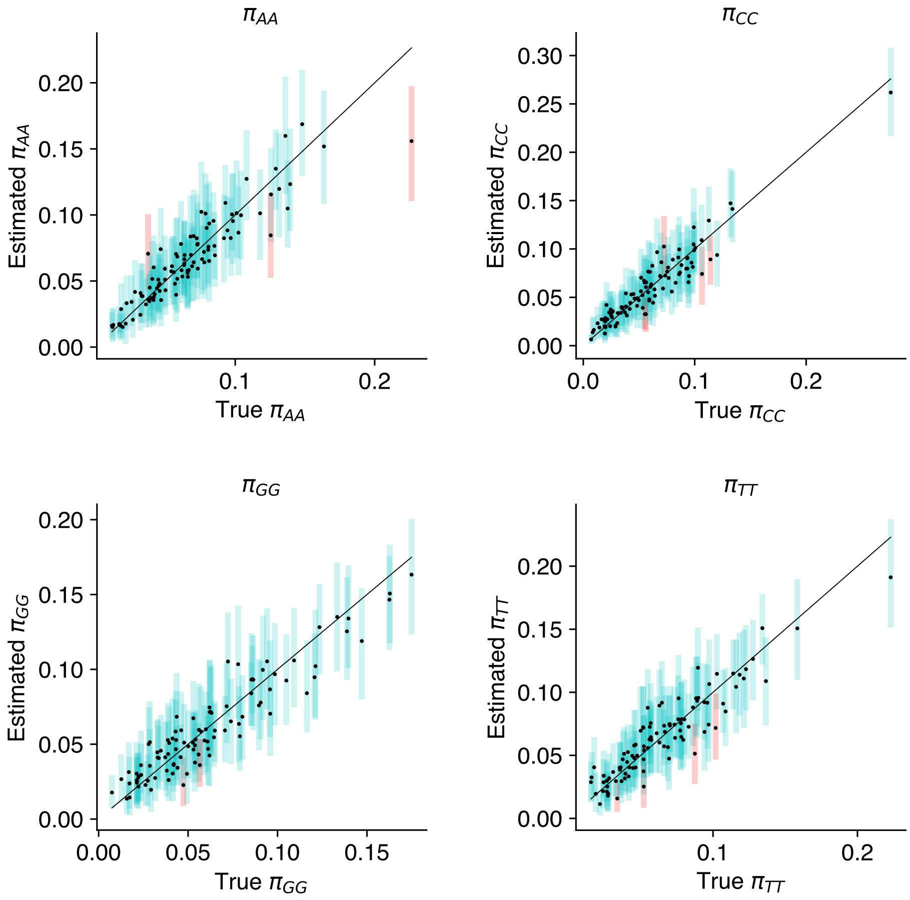

**Figure S5**: Simulation 3: Estimated substitution model frequencies of homozygous genotypes for phased diploid nucleotide data. True vs. estimated frequencies for unphased diploid nucleotide data $\pi_{AA}, \pi_{CC}, \pi_{GG}, \pi_{TT}$ for the GT16 substitution model. Estimated means are shown as points, with true values along the diagonal. The estimated 95% HPD are shown as bars; blue indicates the true value lies within the estimated interval, and red indicates the true value is outside of the estimated interval.

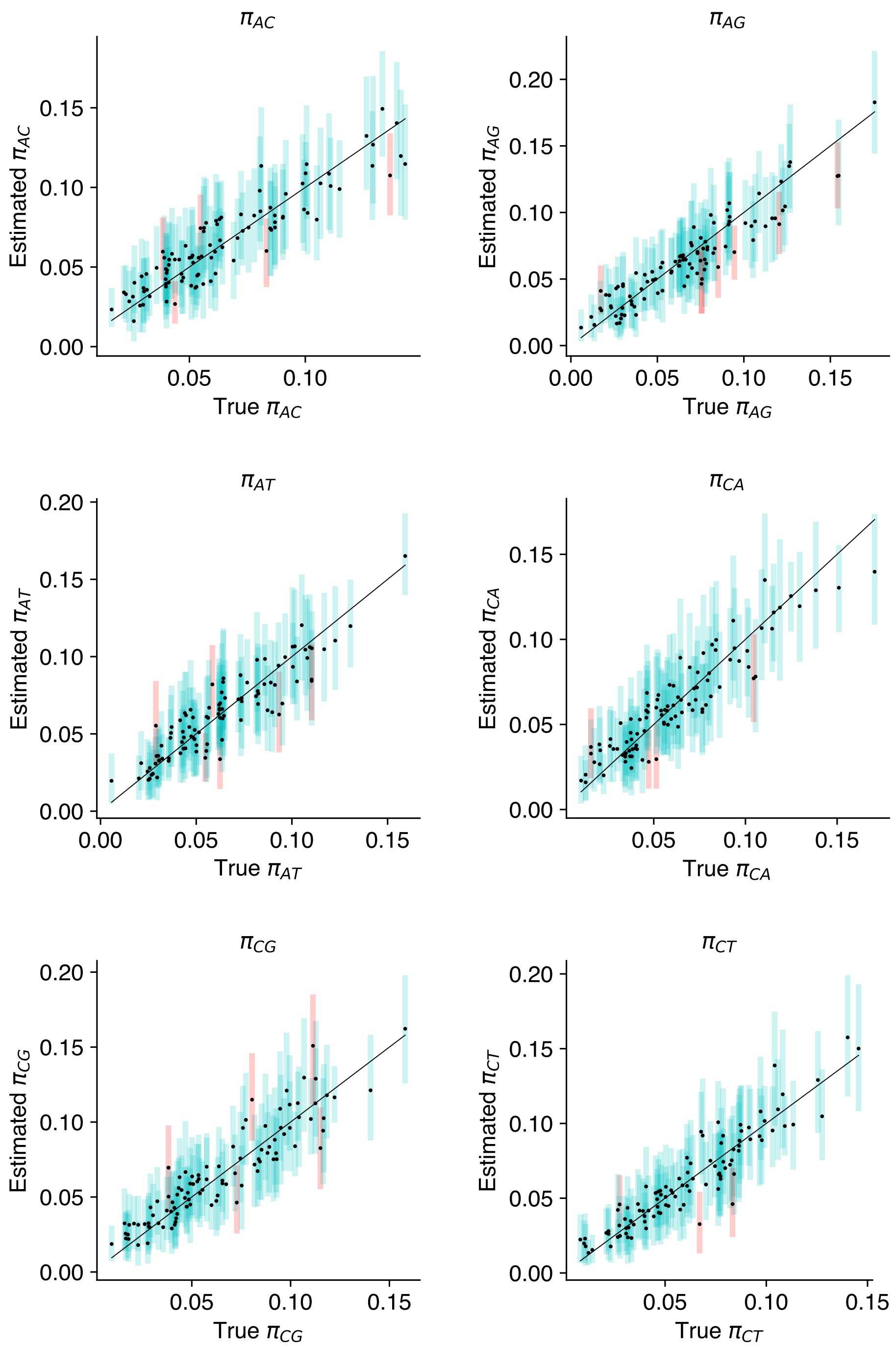

**Figure S6**: Simulation 3: Estimated substitution model frequencies of heterozygous genotypes for phased diploid nucleotide data. True vs. estimated frequencies for heterozygous frequencies in the GT16 substitution model. Estimated means are shown as points, with true values along the diagonal. The estimated 95% HPD are shown as bars; blue indicates the true value lies within the estimated interval, and red indicates the true value is outside of the estimated interval.

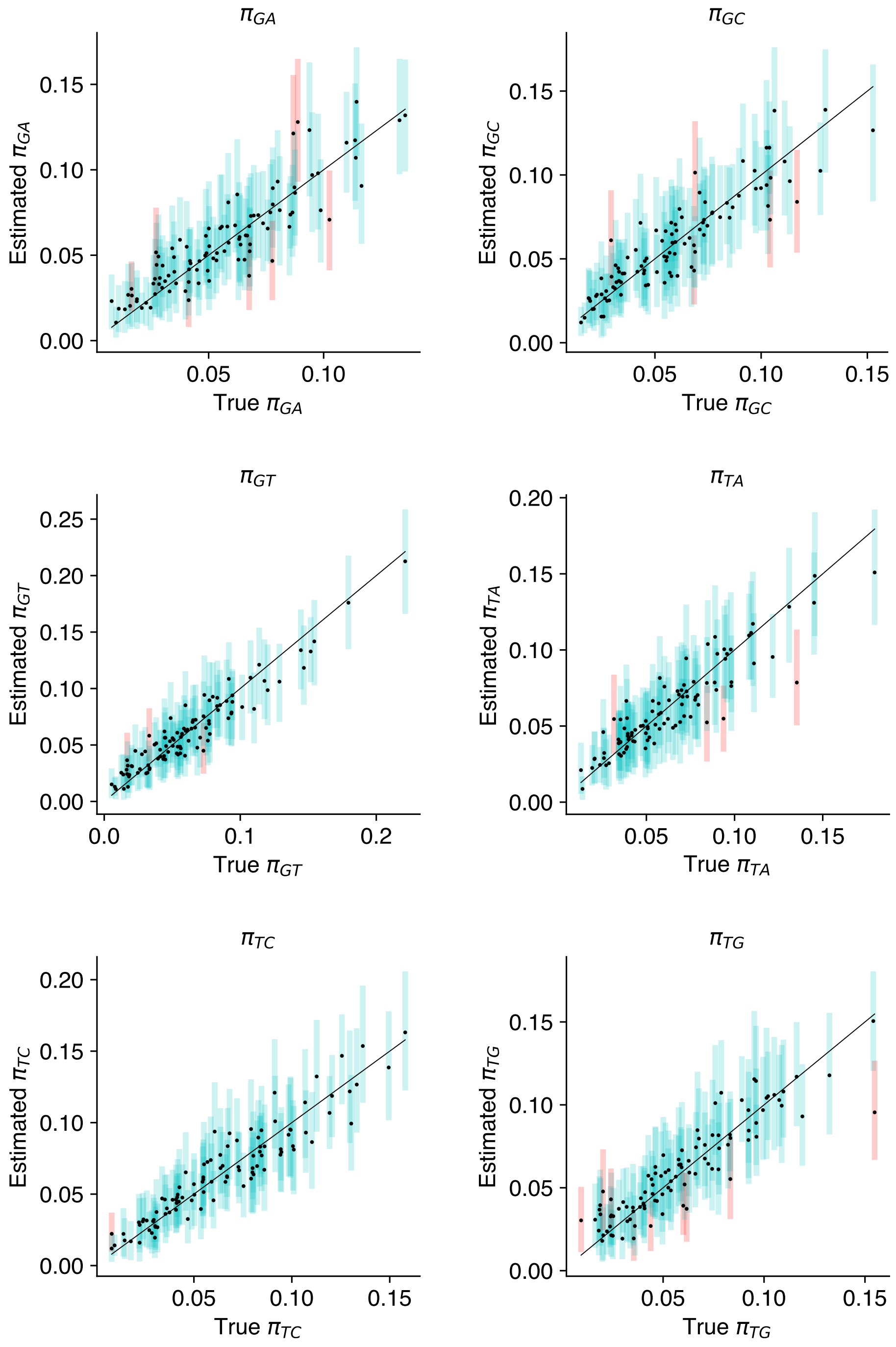

**Figure S7**: Simulation 3: Estimated substitution model frequencies of heterozygous genotypes for phased diploid nucleotide data. True vs. estimated frequencies for heterozygous frequencies in the GT16 substitution model. The estimated 95% HPD are shown as bars; blue indicates the true value lies within the estimated interval, and red indicates the true value is outside of the estimated interval.

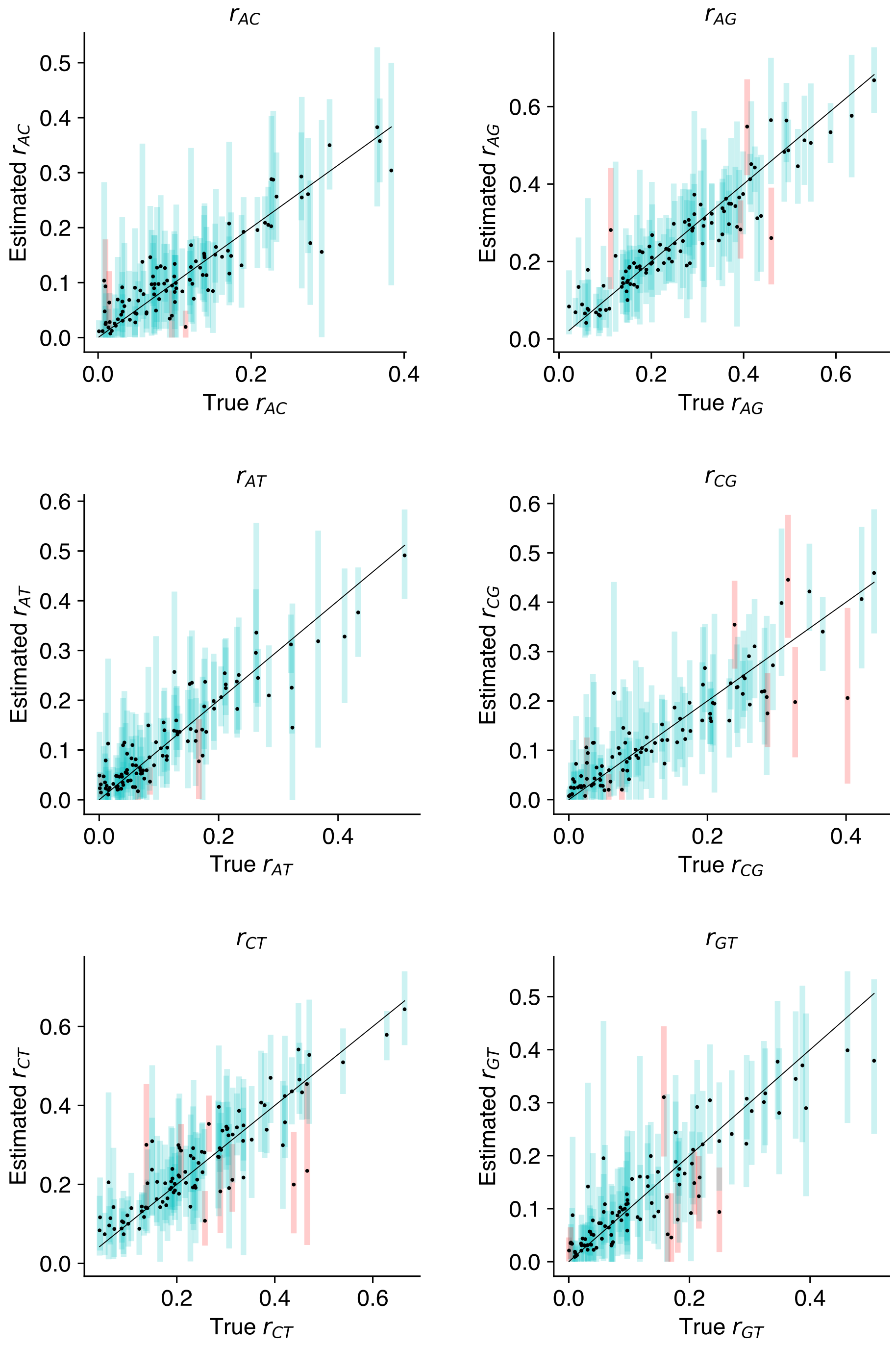

**Figure S8**: Simulation 3: Estimated substitution model relative rates for phased diploid nucleotide data. True vs. estimated relative rates for the GT16 substitution model. The estimated 95% HPD are shown as bars; blue indicates the true value lies within the estimated interval, and red indicates the true value is outside of the estimated interval.

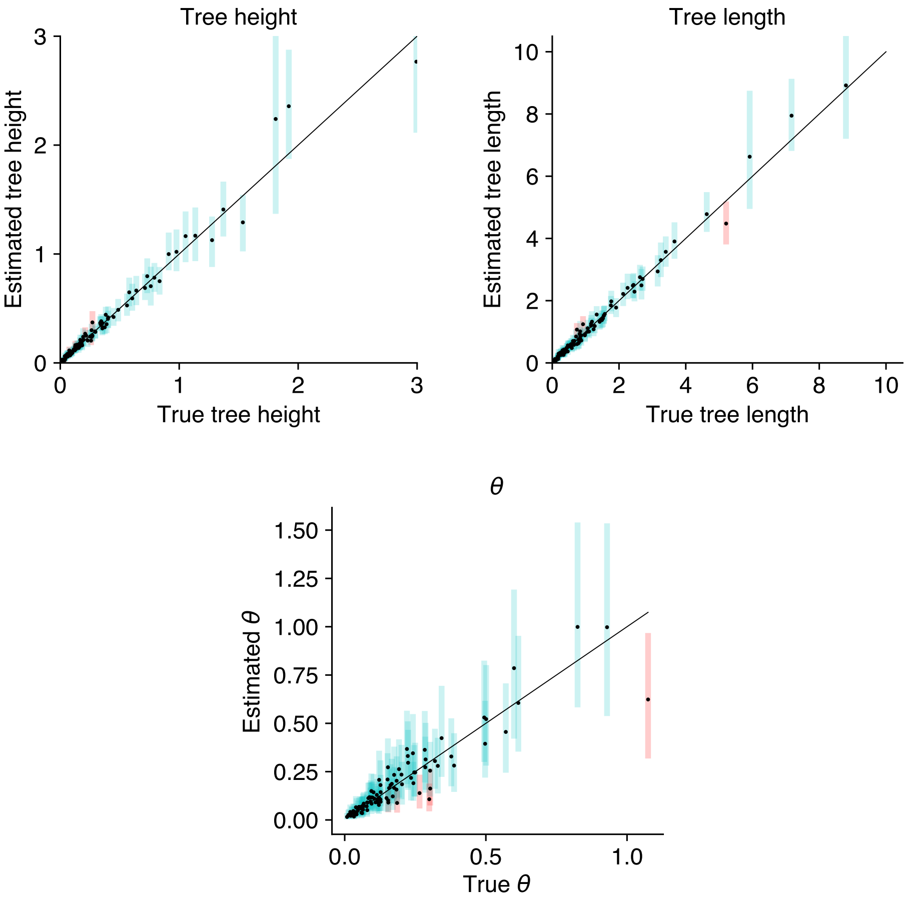

**Figure S9**: Simulation 3: Estimated tree height, tree length and coalescent population size for phased diploid nucleotide data. True vs. estimated values for tree height, tree length and coalescent population size $\theta$. Estimated means are shown as points, with true values along the diagonal. The estimated 95% HPD are shown as bars; blue indicates the true value lies within the estimated interval, and red indicates the true value is outside of the estimated interval.

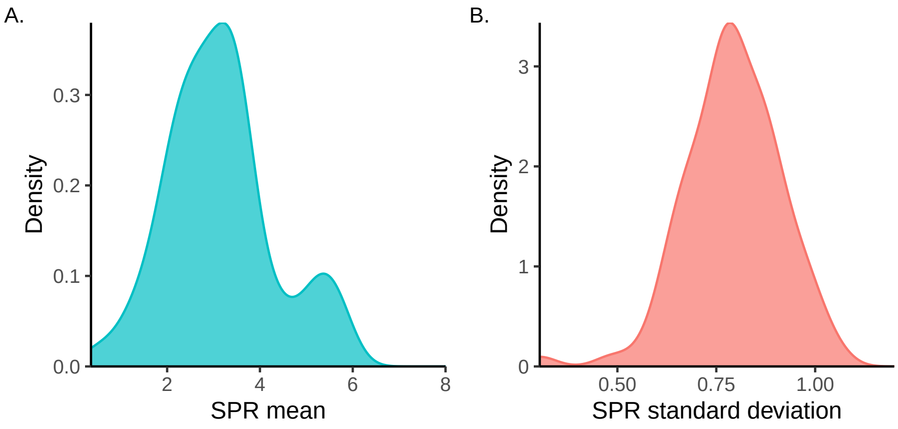

**Figure S10**: Simulation 3: Summary of subtree prune and regraft (SPR) metrics for trees estimated from phased nucleotide data. From left to right, (A) the mean SPR distance between estimated trees and the true tree, (B) the standard deviation of the SPR distance between estimated trees and the true tree.

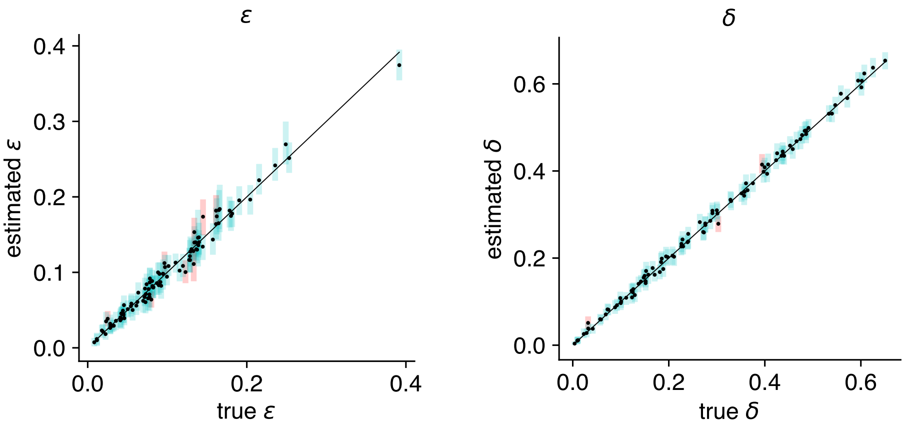

**Figure S11**: Simulation 4: Estimated error parameters for unphased diploid nucleotide data. True vs. estimated error probabilities $\epsilon$ and $\delta$ for the GT16 error model using unphased data. Estimated means are shown as points, with true values along the diagonal. The estimated 95% HPD are shown as bars; blue indicates the true value falls within the estimated interval, and red indicates the true value is outside of the estimated interval.

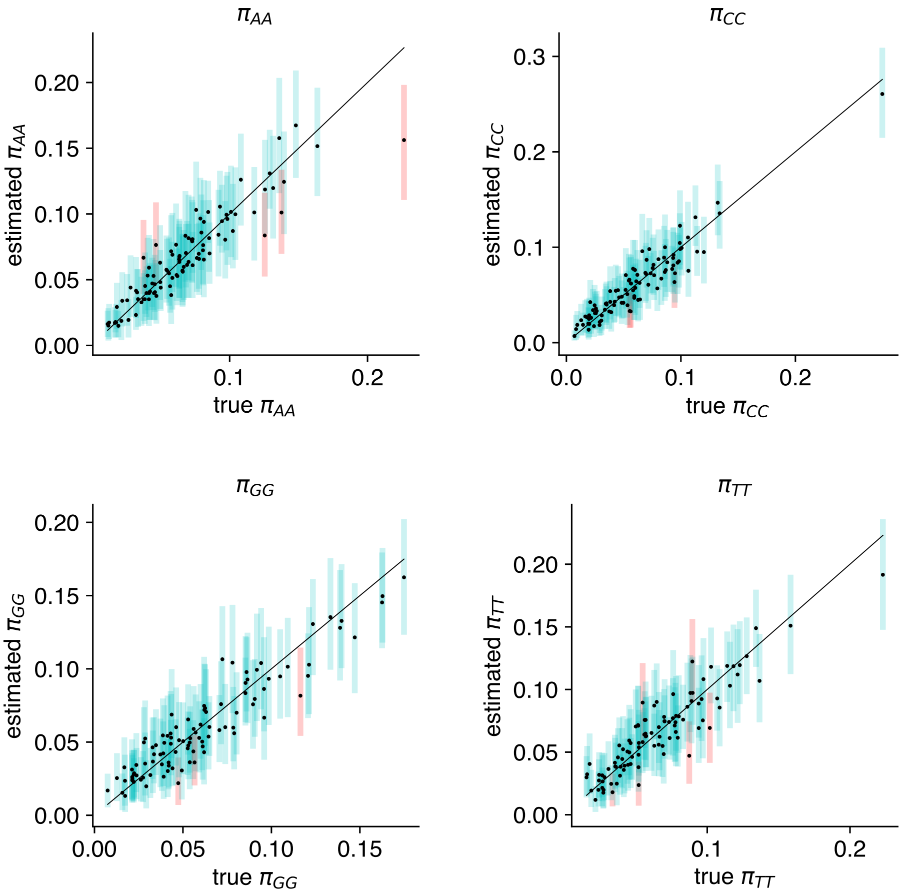

**Figure S12**: Simulation 4: Estimated substitution model frequencies of homozygous genotypes for unphased diploid nucleotide data. True vs. estimated frequencies for unphased diploid nucleotide data $\pi_{AA}, \pi_{CC}, \pi_{GG}, \pi_{TT}$ for the GT16 substitution model. Estimated means are shown as points, with true values along the diagonal. The estimated 95% HPD are shown as bars; blue indicates the true value lies within the estimated interval, and red indicates the true value is outside of the estimated interval.

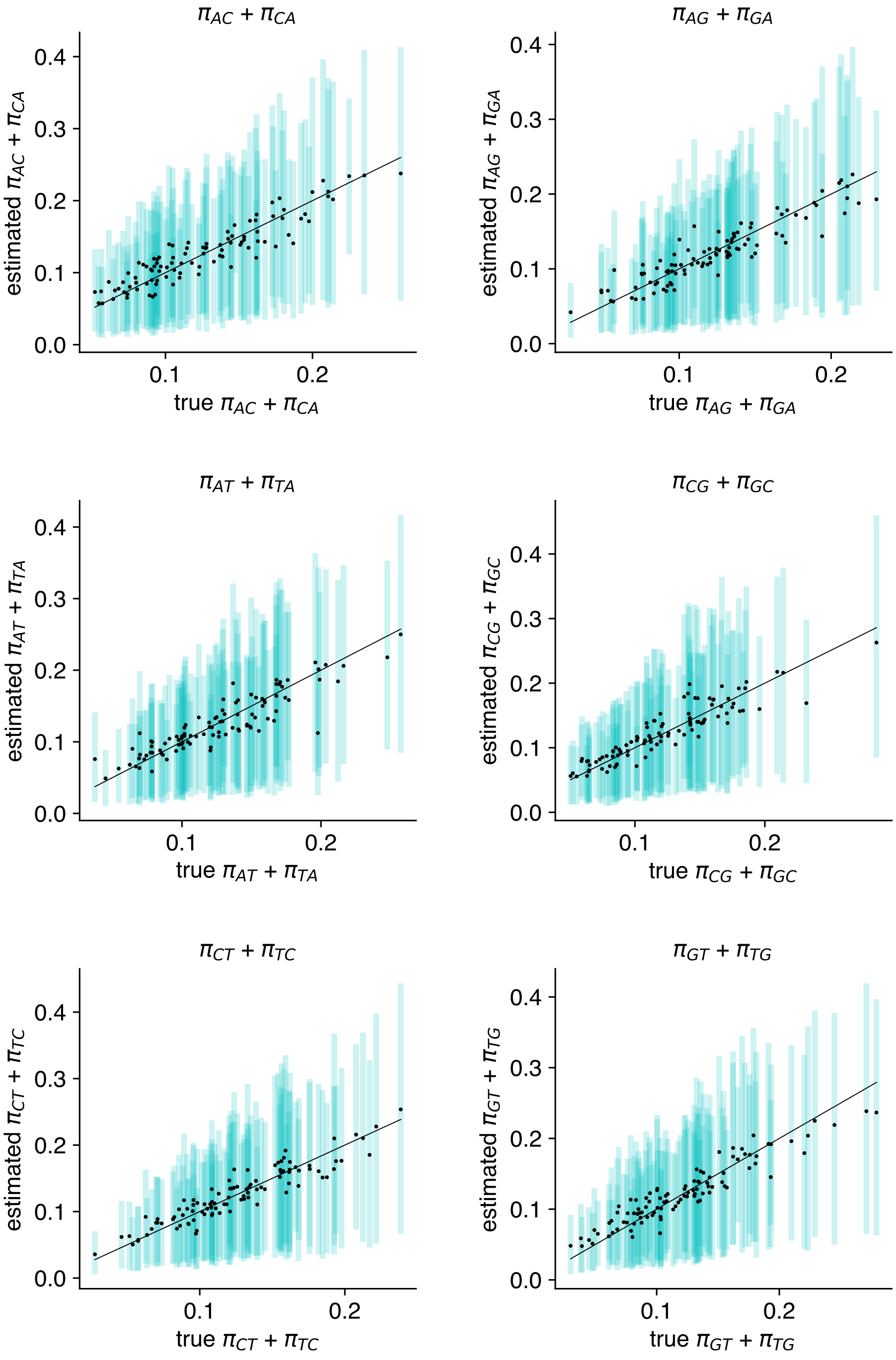

**Figure S13**: Simulation 4: Estimated substitution model frequencies of heterozygous genotypes for unphased diploid nucleotide data. True vs. estimated frequencies for the sum of heterozygous pairs $\pi_{ij} + \pi_{ji},\ i \neq j$ in the GT16 substitution model. Estimated means are shown as points, with true values along the diagonal. The estimated 95% HPD are shown as bars; blue indicates the true value lies within the estimated interval, and red indicates the true value is outside of the estimated interval.

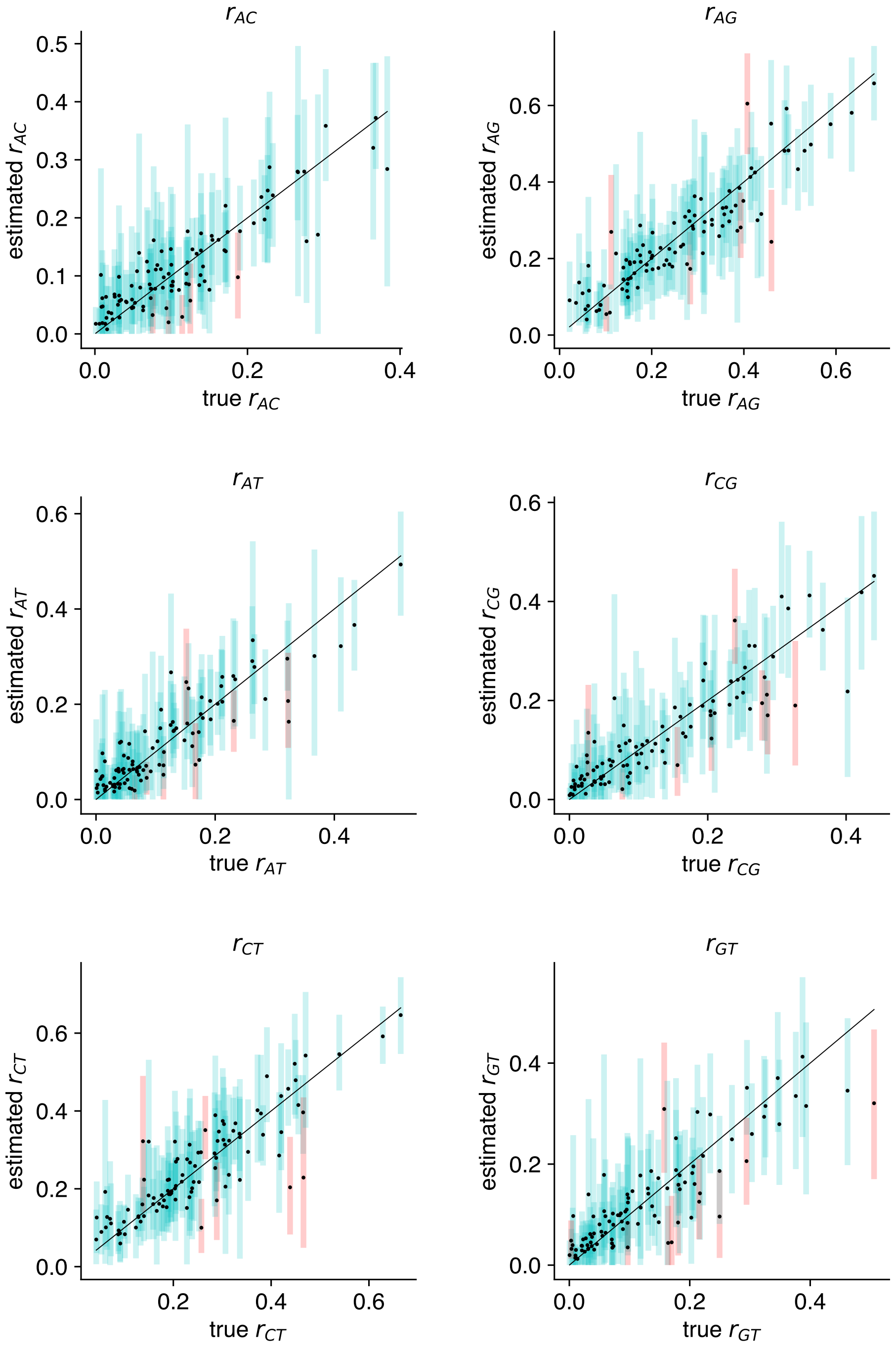

**Figure S14**: Simulation 4: Estimated substitution model relative rates for unphased diploid nucleotide data. True vs. estimated relative rates for the GT16 substitution model. Estimated means are shown as points, with true values along the diagonal. The estimated 95% HPD are shown as bars; blue indicates the true value lies within the estimated interval, and red indicates the true value is outside of the estimated interval.

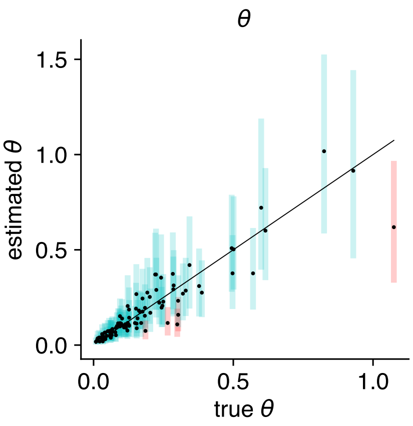

**Figure S15**: Simulation 4: Estimated coalescent population size for unphased diploid nucleotide data. True vs. estimated $\theta$ for the coalescent model. Estimated means are shown as points, with true values along the diagonal. The estimated 95% HPD are shown as bars; blue indicates the true value lies within the estimated interval, and red indicates the true value is outside of the estimated interval.

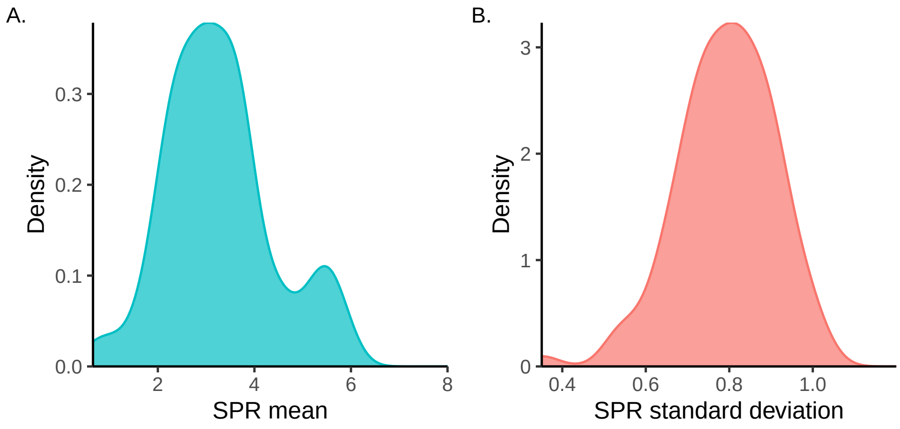

**Figure S16**: Simulation 4: Summary of subtree prune and regraft (SPR) metrics for trees estimated from unphased nucleotide data. From left to right, (A) the mean SPR distance between estimated trees and the true tree, (B) the standard deviation of the SPR distance between estimated trees and the true tree.

**Table S2**: Simulation 5: Coverage comparison for model parameters with and without the GT16 error model.

| Parameter | 95% HPD coverage with error model | 95% HPD coverage without error model |
|---:|:---:|:---:|
| $\delta$ | 95% | - |
| $\epsilon$ | 92% | - |
| $\pi_{AA}$ | 97% | 9% |
| $\pi_{AC}$ | 95% | 23% |
| $\pi_{AG}$ | 93% | 21% |
| $\pi_{AT}$ | 94% | 15% |
| $\pi_{CA}$ | 96% | 22% |
| $\pi_{CC}$ | 95% | 4% |
| $\pi_{CG}$ | 95% | 18% |
| $\pi_{CT}$ | 97% | 20% |
| $\pi_{GA}$ | 92% | 25% |
| $\pi_{GC}$ | 95% | 23% |
| $\pi_{GG}$ | 98% | 10% |
| $\pi_{GT}$ | 97% | 15% |
| $\pi_{TA}$ | 96% | 18% |
| $\pi_{TC}$ | 99% | 18% |
| $\pi_{TG}$ | 91% | 21% |
| $\pi_{TT}$ | 96% | 8% |
| $r_{AC}$ | 96% | 19% |
| $r_{AG}$ | 96% | 22% |
| $r_{AT}$ | 96% | 20% |
| $r_{CG}$ | 91% | 15% |
| $r_{CT}$ | 92% | 25% |
| $r_{GT}$ | 91% | 16% |
| $\theta$ | 94% | 0% |
| Tree height | 95% | 10% |
| Tree length | 92% | 0% |

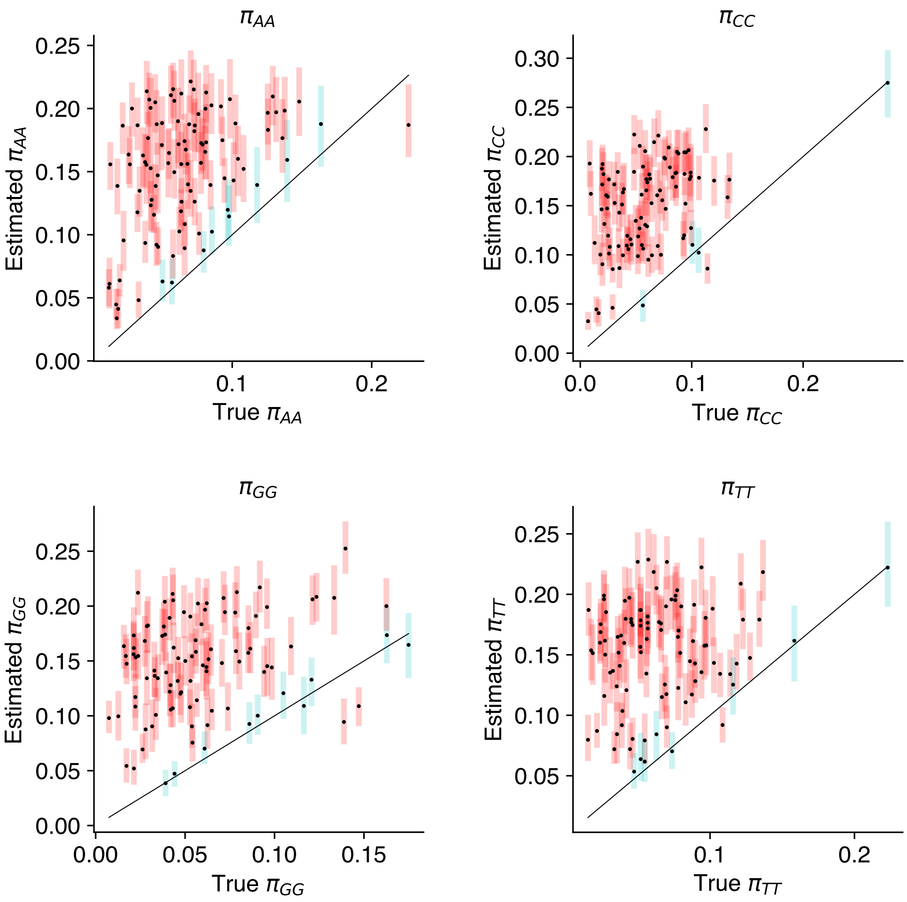

**Figure S17**: Simulation 5: Estimated substitution model frequencies of homozygous genotypes for phased diploid nucleotide data without an error model. True vs. estimated frequencies for unphased diploid nucleotide data $\pi_{AA}, \pi_{CC}, \pi_{GG}, \pi_{TT}$ for the GT16 substitution model. Estimated means are shown as points, with true values along the diagonal. The estimated 95% HPD are shown as bars; blue indicates the true value lies within the estimated interval, and red indicates the true value is outside of the estimated interval.

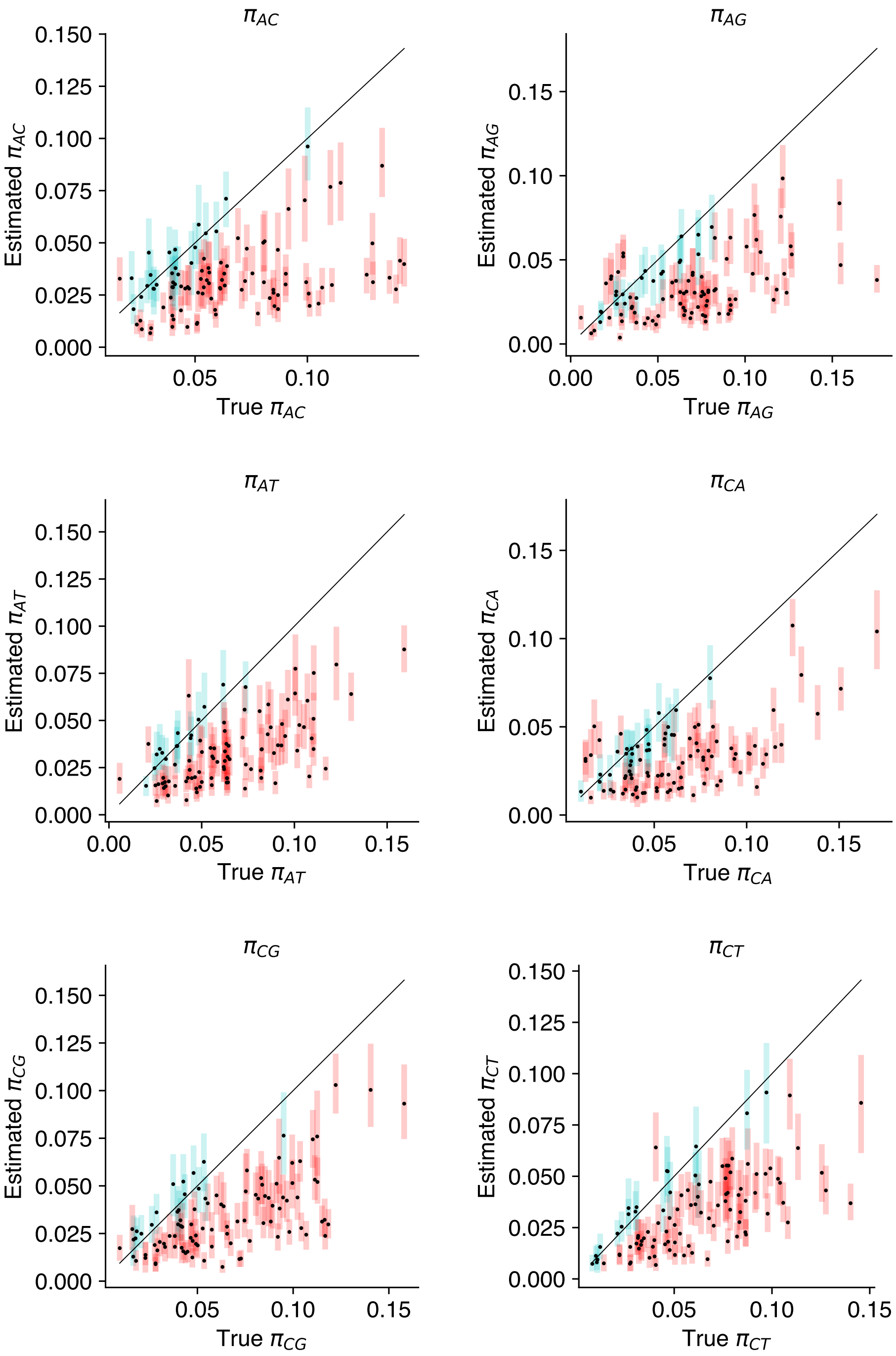

**Figure S18**: Simulation 5: Estimated substitution model frequencies of heterozygous genotypes for phased diploid nucleotide data without an error model. True vs. estimated frequencies for heterozygous frequencies in the GT16 substitution model. Estimated means are shown as points, with true values along the diagonal. The estimated 95% HPD are shown as bars; blue indicates the true value lies within the estimated interval, and red indicates the true value is outside of the estimated interval.

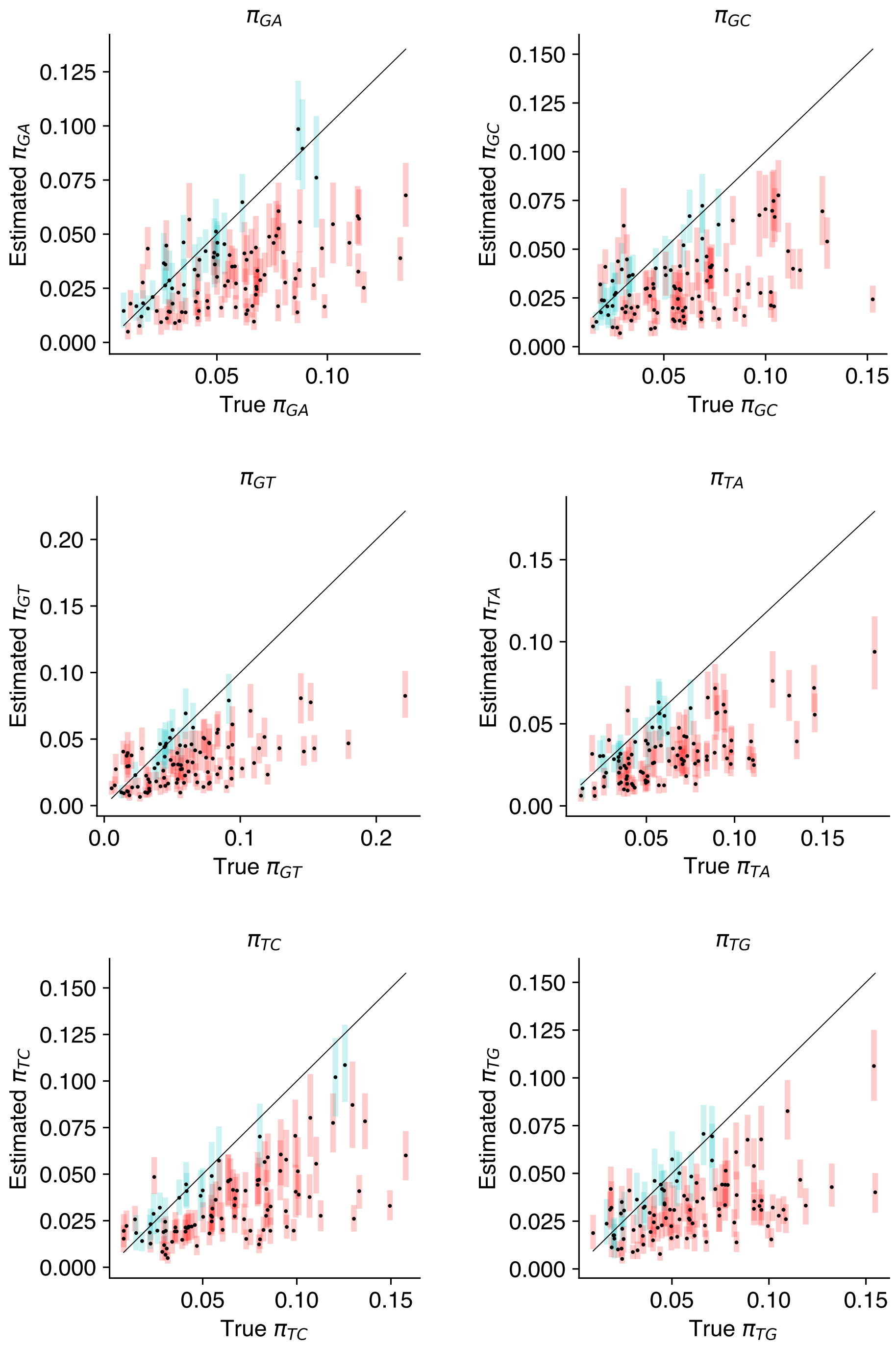

**Figure S19**: Simulation 5: Estimated substitution model frequencies of heterozygous genotypes for phased diploid nucleotide data without an error model. True vs. estimated frequencies for heterozygous frequencies in the GT16 substitution model. The estimated 95% HPD are shown as bars; blue indicates the true value lies within the estimated interval, and red indicates the true value is outside of the estimated interval.

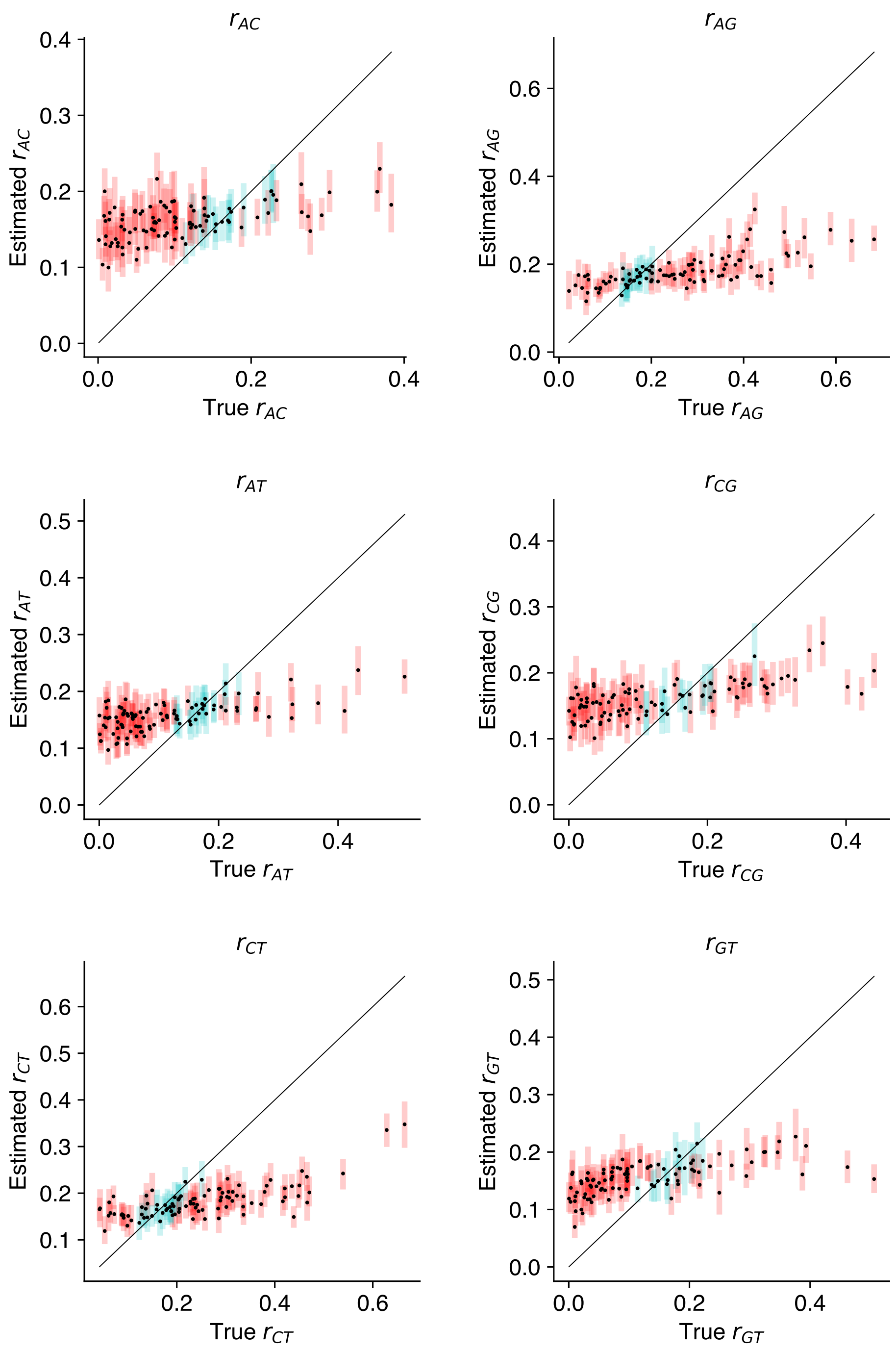

**Figure S20**: Simulation 5: Estimated substitution model relative rates for phased diploid nucleotide data without an error model. True vs. estimated relative rates for the GT16 substitution model. The estimated 95% HPD are shown as bars; blue indicates the true value lies within the estimated interval, and red indicates the true value is outside of the estimated interval.

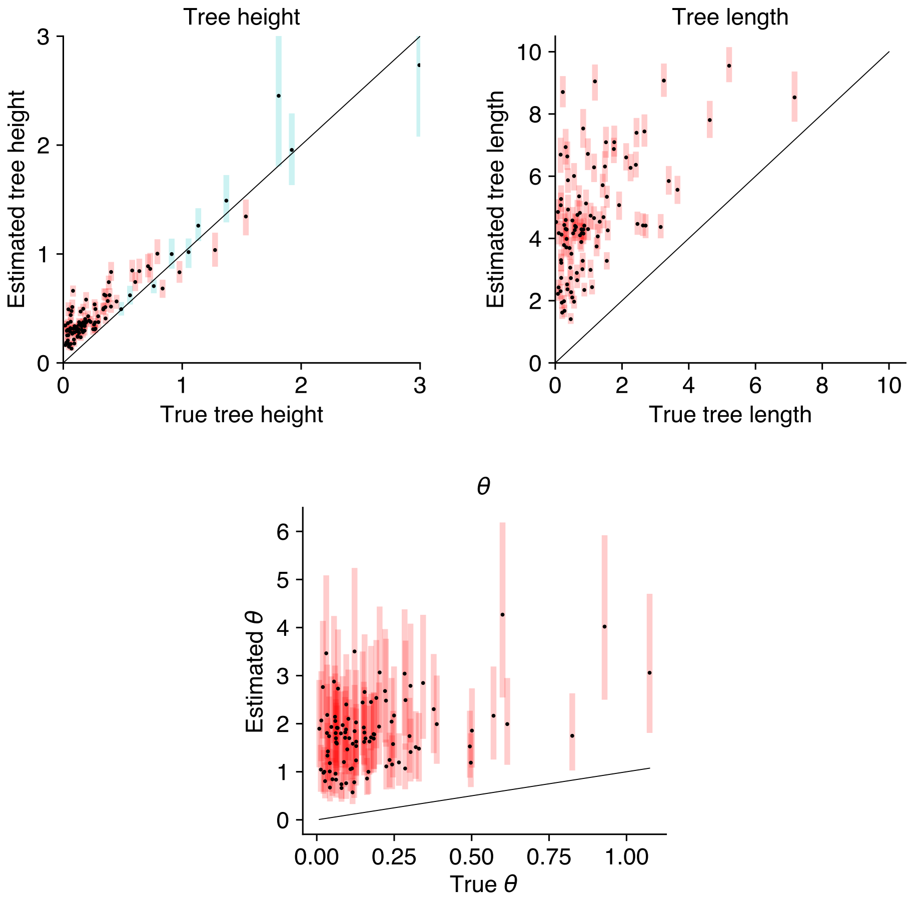

**Figure S21**: Simulation 5: Estimated tree height, tree length and coalescent population size for phased diploid nucleotide data without an error model. True vs. estimated values for tree height, tree length and coalescent population size $\theta$. Estimated means are shown as points, with true values along the diagonal. The estimated 95% HPD are shown as bars; blue indicates the true value lies within the estimated interval, and red indicates the true value is outside of the estimated interval.

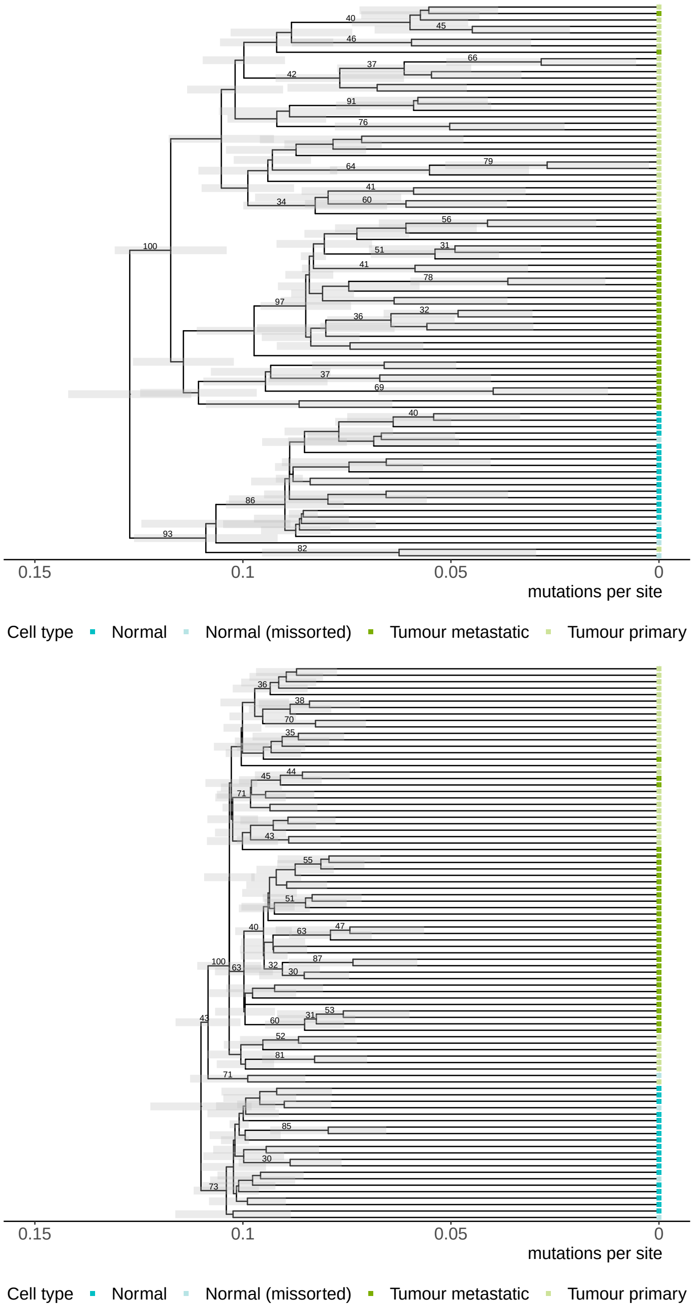

**Figure S22**: Maximium clade credibility trees for a colorectal cancer patient using the GT16 model with the normal cells as the outgroup. Trees were estimated with an error model (top) and without an error model (bottom). Cells are colored by their cell types: normal cells (dark blue), normal cells missorted during data collection (light blue), the metastatic tumour from the liver (dark green), and primary tumour from the colon (light green). The posterior clade support for clades with greater than 30% support are shown on the branches.

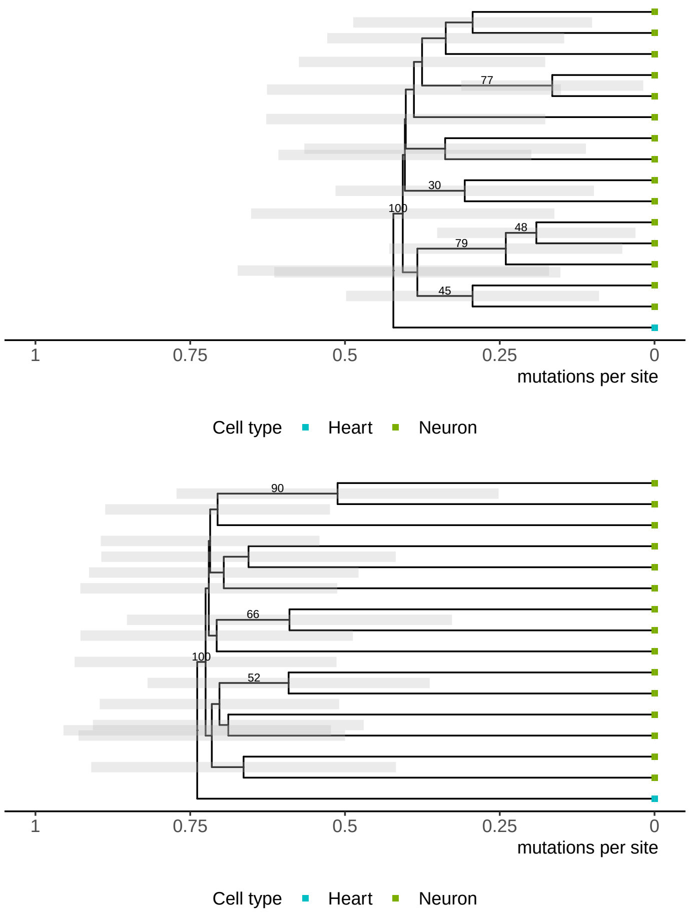

**Figure S23**: Maximum clade credibility trees for a healthy patient using the GT16 model with the heart cell as the outgroup. Trees were estimated with an error model (top) and without an error model (bottom). Each cell is colored by its cell type: blood cell (blue) and neuron cells (green). The posterior clade support for clades with greater than 30% support are shown on the branches.

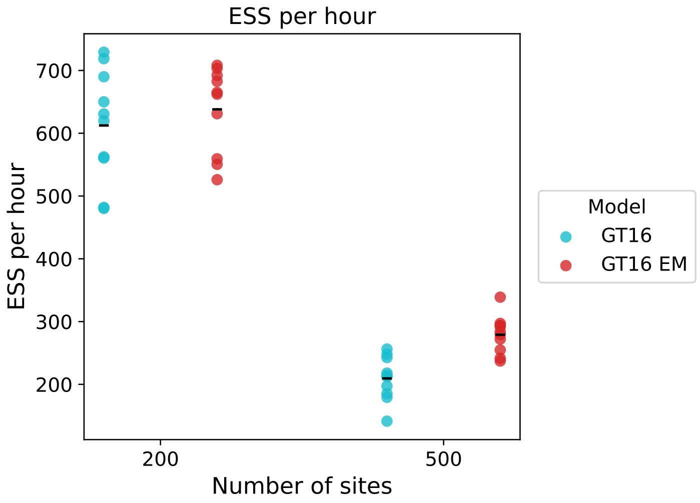

**Figure S24**: Simulation 6: Convergence comparison of the GT16 error model (EM) with the baseline non-error GT16 model showing the minumum estimated sample size (ESS) divided by the total runtime for each experimental replicate. Ten experimental replicates were performed for each setting. The average ESS per hour is indicated by a horizontal bar.

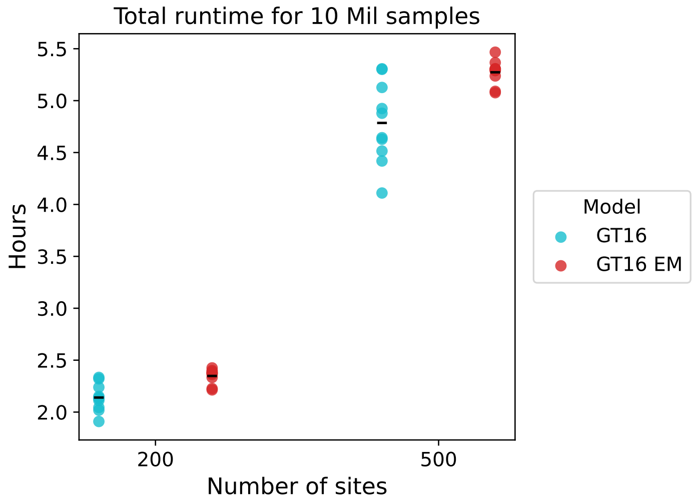

**Figure S25**: Simulation 6: Timing comparison of the GT16 error model (EM) with the baseline non-error GT16 model showing runtime in hours required to produce 10 million samples. Ten experimental replicates were performed for each setting. The average runtime is indicated by a horizontal bar.

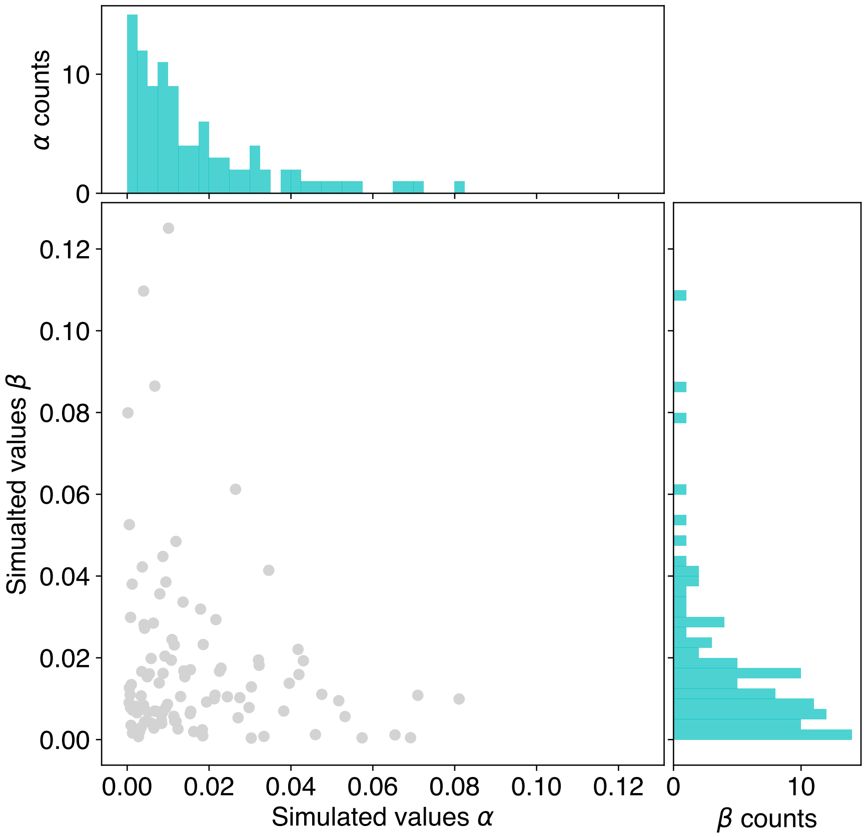

**Figure S26**: Simulated values for false positive ($\alpha$) and false negative errors ($\beta$) from simulations 1 and 2. The joint distribution of the two simulated error parameters are shown by the scatter plot. The individual distributions for each parameter are shown by the histograms.

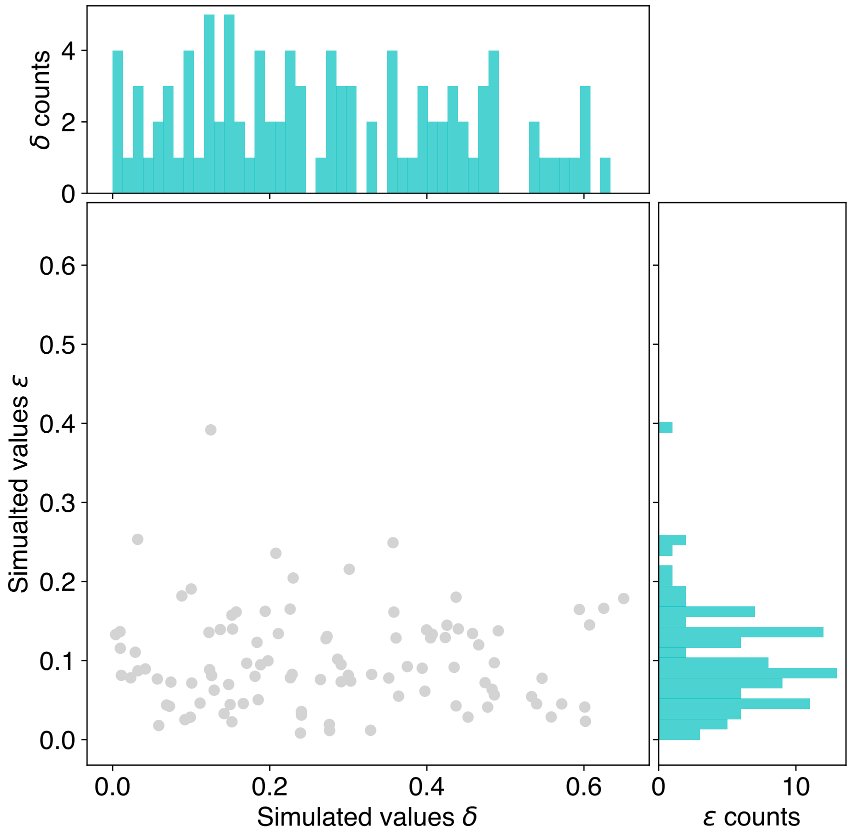

**Figure S27**: Simulated values for allelic dropout ($\delta$) and the combined amplification and sequencing error ($\epsilon$) from simulations 3-5. The joint distribution of the two simulated error parameters are shown by the scatter plot. The individual distributions for each parameter are shown by the histograms.

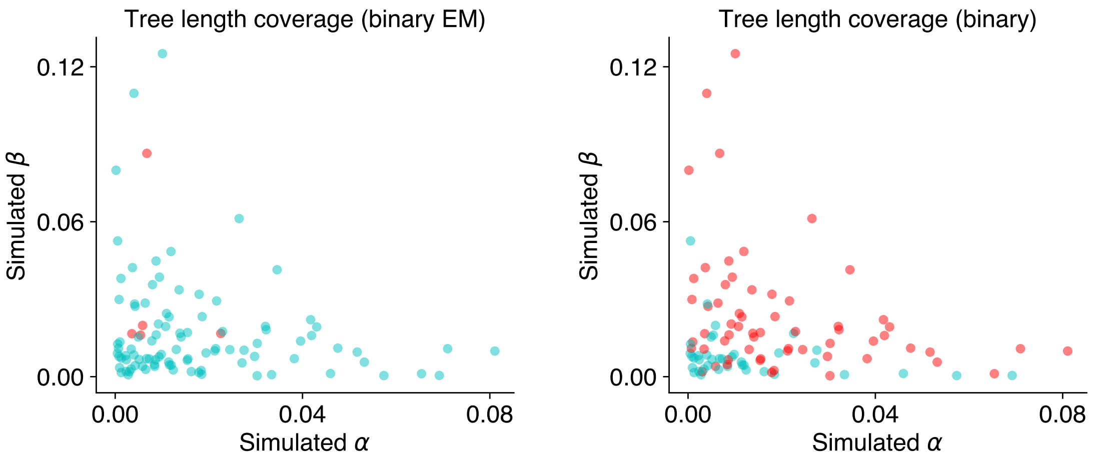

**Figure S28**: Tree length coverage for varying levels of error using the binary error model (EM) and the binary model without error. Simulated values for false positive ($\alpha$) and false negative errors ($\beta$) are shown on the two axis. Each point represents a randomly simulated dataset; blue indicates the estimated interval covers the true tree length and red indicates otherwise.

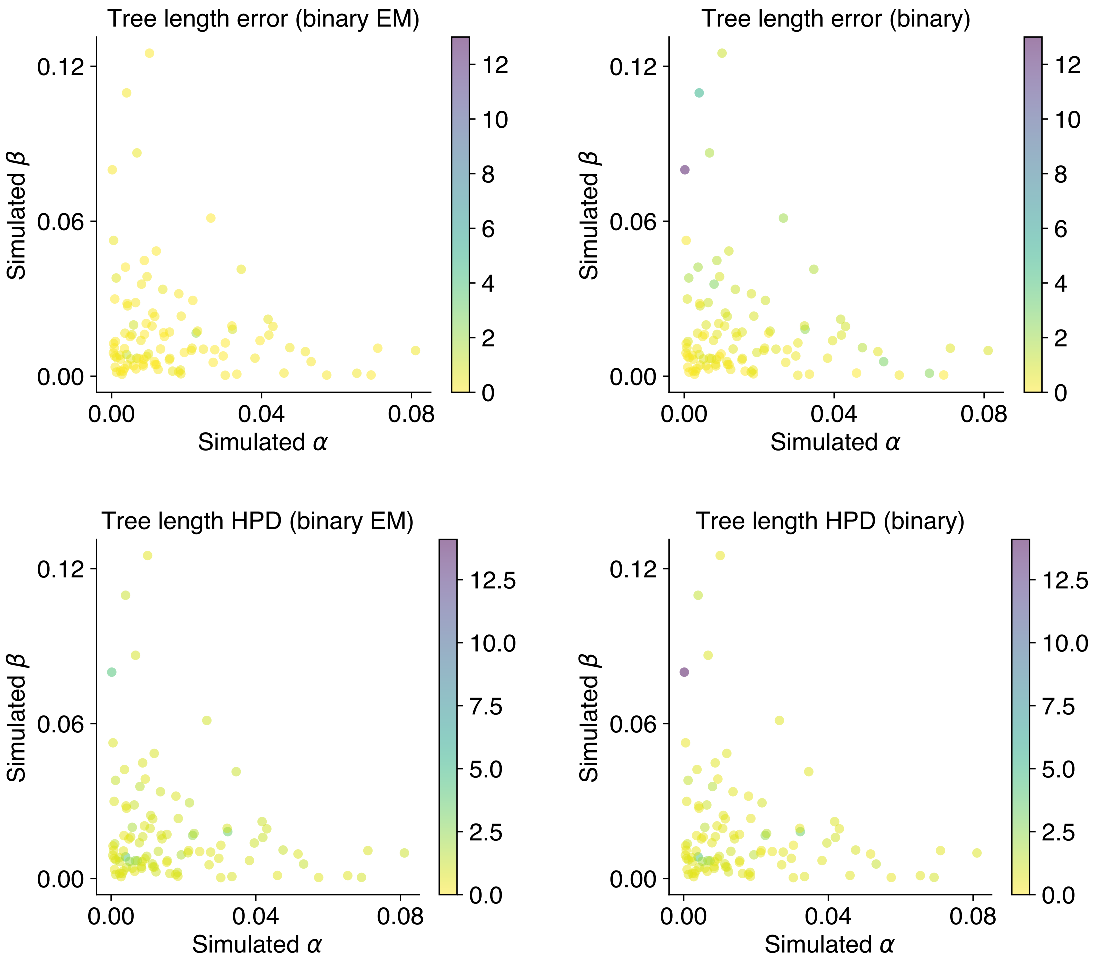

**Figure S29**: Tree length estimates for varying error levels using the binary error model (EM) and the binary model without error. Simulated values for false positive ($\alpha$) and false negative errors ($\beta$) are shown on the two axis. The top plots show the magnitude of error, which is calculated as the difference between the estimated mean tree length and the true tree length. The size of the estimated HPD interval is shown on the bottom plots.

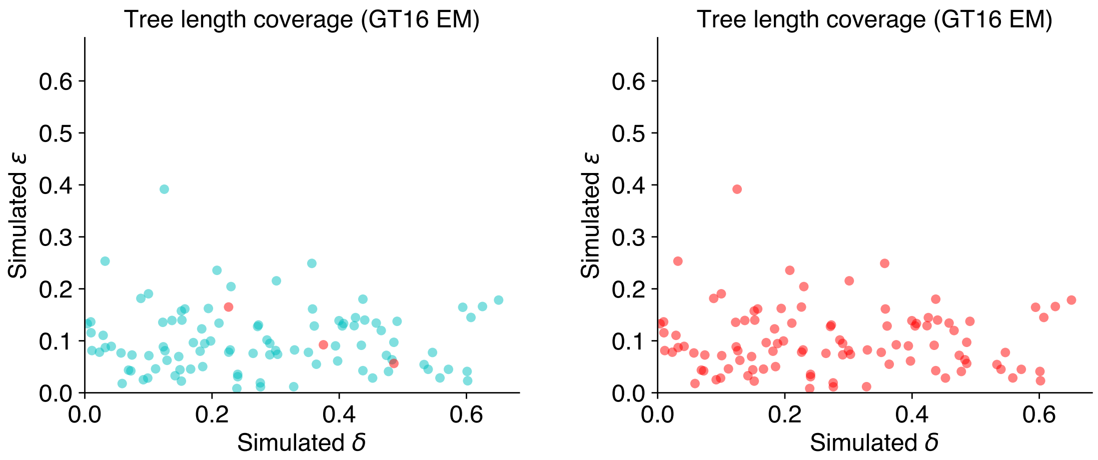

**Figure S30**: Tree length coverage for varying levels of error using the GT16 error model (EM) and the GT16 model without error. Simulated values for allelic dropout ($\delta$) and the combined amplification and sequencing error ($\epsilon$) are shown on the two axis. Each point represents a randomly simulated dataset; blue indicates the estimated interval covers the true tree length and red indicates otherwise.

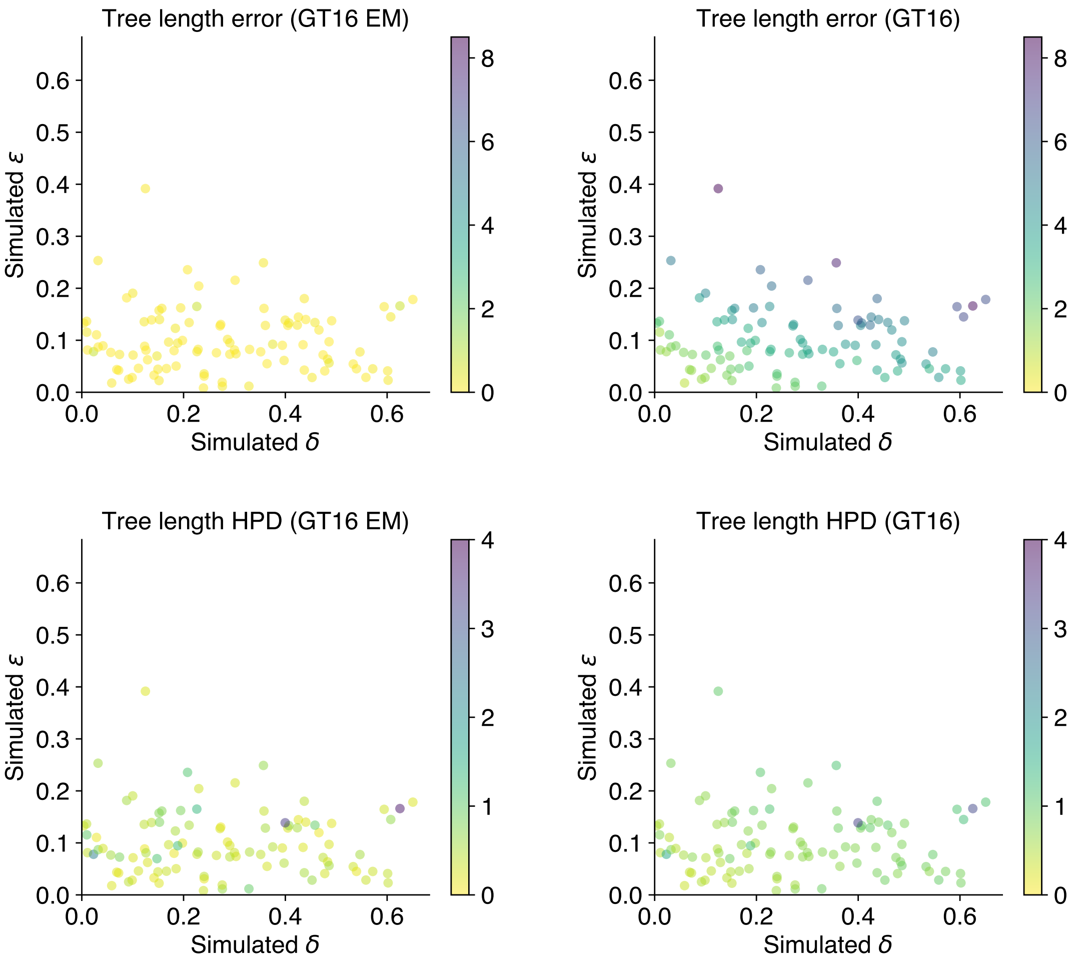

**Figure S31**: Tree length estimates for varying levels of error using the GT16 error model (EM) and the GT16 model without error. Simulated values for allelic dropout ($\delta$) and the combined amplification and sequencing error ($\epsilon$) are shown on the two axis. The top plots show the magnitude of error, which is calculated as the difference between the estimated mean tree length and the true tree length. The size of the estimated HPD intervals is shown on the bottom plots.

**Table S3**: Simulation 7a: Parameter coverage (95% HPD) for the GT16 error model on datasets with varying levels of allelic dropout error $\delta \in [0.1, 0.25, 0.5, 0.8]$ and amplification/sequencing error $\epsilon = 0.001$. Each setup was repeated 10 times, the coverage statistics for each model parameter are summarized below.

| | True $\delta$ | | | |
|---|---|---|---|---|
| Parameter coverage | 0.1 | 0.25 | 0.5 | 0.8 |
| $\delta$ | 90% | 80% | 90% | 90% |
| $\epsilon$ | 90% | 80% | 100% | 60% |
| $\pi_{AA}$ | 100% | 90% | 100% | 100% |
| $\pi_{AC}$ | 100% | 100% | 90% | 90% |
| $\pi_{AG}$ | 100% | 90% | 90% | 100% |
| $\pi_{AT}$ | 100% | 100% | 100% | 90% |
| $\pi_{CA}$ | 90% | 100% | 100% | 90% |
| $\pi_{CC}$ | 80% | 90% | 100% | 100% |
| $\pi_{CG}$ | 100% | 100% | 90% | 100% |
| $\pi_{CT}$ | 90% | 90% | 100% | 100% |
| $\pi_{GA}$ | 90% | 100% | 100% | 100% |
| $\pi_{GC}$ | 100% | 80% | 100% | 90% |
| $\pi_{GG}$ | 100% | 100% | 80% | 90% |
| $\pi_{GT}$ | 100% | 80% | 90% | 100% |
| $\pi_{TA}$ | 100% | 90% | 90% | 90% |
| $\pi_{TC}$ | 100% | 100% | 100% | 100% |
| $\pi_{TG}$ | 70% | 100% | 90% | 100% |
| $\pi_{TT}$ | 90% | 100% | 100% | 100% |
| $r_{AC}$ | 100% | 100% | 100% | 100% |
| $r_{AG}$ | 90% | 80% | 100% | 100% |
| $r_{AT}$ | 100% | 90% | 100% | 90% |
| $r_{CG}$ | 100% | 90% | 100% | 100% |
| $r_{CT}$ | 100% | 90% | 100% | 100% |
| $r_{GT}$ | 100% | 100% | 100% | 90% |
| $\theta$ | 100% | 100% | 100% | 100% |
| Tree height | 100% | 100% | 90% | 100% |
| Tree length | 100% | 90% | 100% | 90% |

**Table S4**: Simulation 7b: Parameter coverage (95% HPD) for the GT16 error model on datasets with varying levels of amplification/sequencing error $\epsilon \in [0.001, 0.01, 0.05, 0.1]$ and allelic dropout error $\delta = 0.5$. Each setup was repeated 10 times, the coverage statistics for each model parameter are summarized below.

| | True $\epsilon$ | | | |
|---|---|---|---|---|
| Parameter coverage | 0.001 | 0.01 | 0.05 | 0.1 |
| $\delta$ | 80% | 90% | 70% | 100% |
| $\epsilon$ | 80% | 100% | 80% | 90% |
| $\pi_{AA}$ | 100% | 100% | 100% | 100% |
| $\pi_{AC}$ | 100% | 100% | 70% | 100% |
| $\pi_{AG}$ | 100% | 90% | 80% | 100% |
| $\pi_{AT}$ | 100% | 100% | 80% | 100% |
| $\pi_{CA}$ | 80% | 100% | 100% | 100% |
| $\pi_{CC}$ | 90% | 90% | 80% | 100% |
| $\pi_{CG}$ | 100% | 100% | 90% | 100% |
| $\pi_{CT}$ | 100% | 90% | 100% | 100% |
| $\pi_{GA}$ | 80% | 90% | 100% | 90% |
| $\pi_{GC}$ | 80% | 100% | 90% | 100% |
| $\pi_{GG}$ | 100% | 100% | 100% | 80% |
| $\pi_{GT}$ | 100% | 100% | 100% | 100% |
| $\pi_{TA}$ | 90% | 80% | 90% | 100% |
| $\pi_{TC}$ | 90% | 90% | 90% | 100% |
| $\pi_{TG}$ | 100% | 100% | 100% | 90% |
| $\pi_{TT}$ | 100% | 100% | 80% | 100% |
| $r_{AC}$ | 100% | 90% | 100% | 100% |
| $r_{AG}$ | 100% | 100% | 100% | 100% |
| $r_{AT}$ | 100% | 80% | 90% | 100% |
| $r_{CG}$ | 100% | 100% | 100% | 100% |
| $r_{CT}$ | 100% | 100% | 100% | 100% |
| $r_{GT}$ | 100% | 100% | 100% | 100% |
| $\theta$ | 90% | 100% | 100% | 100% |
| Tree height | 100% | 100% | 80% | 100% |
| Tree length | 90% | 90% | 80% | 100% |

**Table S5**: Simulation 7c: Parameter coverage (95% HPD) for the binary error model on datasets with varying levels of false positive error $\alpha \in [0.001, 0.01, 0.05, 0.1]$ and false negative error $\beta = 0.5$. Each setup was repeated 10 times, the coverage statistics for each model parameter are summarized below.

| | True $\alpha$ | | | |
|---|---|---|---|---|
| Parameter coverage | 0.001 | 0.01 | 0.05 | 0.1 |
| $\alpha$ | 100% | 100% | 100% | 100% |
| $\beta$ | 100% | 100% | 100% | 100% |
| $\lambda$ | 100% | 100% | 90% | 100% |
| Birthrate | 100% | 100% | 90% | 90% |
| Tree height | 100% | 80% | 100% | 100% |
| Tree length | 100% | 100% | 100% | 100% |

**Table S6**: Simulation 7d: Parameter coverage (95% HPD) for the binary error model on datasets with varying levels of false negative error $\beta \in [0.1, 0.25, 0.5, 0.6]$ and false positive error $\alpha = 0.001$. Each setup was repeated 10 times, the coverage statistics for each model parameter are summarized below.

| | True $\beta$ | | | |
|---|---|---|---|---|
| Parameter coverage | 0.1 | 0.25 | 0.5 | 0.6 |
| $\alpha$ | 90% | 100% | 100% | 100% |
| $\beta$ | 80% | 90% | 100% | 100% |
| $\lambda$ | 100% | 90% | 90% | 100% |
| Birthrate | 80% | 90% | 100% | 100% |
| Tree height | 90% | 90% | 100% | 100% |
| Tree length | 90% | 80% | 100% | 100% |
